# Conserved cholesterol-related activities of Dispatched drive Sonic hedgehog shedding from the cell membrane

**DOI:** 10.1101/2020.10.19.346395

**Authors:** K. Ehring, D. Manikowski, J. Froese, J. Goretzko, P. Jakobs, F. Gude, U. Rescher, K. Grobe

**Author notes:** Correspondence to: K. Grobe.

## Abstract

The Sonic hedgehog (Shh) pathway controls embryonic development and tissue homeostasis after birth. Long-lasting questions about this pathway are how dual-lipidated, firmly plasma membrane-associated Shh ligand is released from producing cells to signal to distant target cells, and how the resistance-nodulation-division transporter Dispatched (Disp) regulates this process. Here we show that Disp inactivation in Shh expressing cells specifically impairs proteolytic Shh release from its lipidated terminal peptides, a process called ectodomain shedding. Shh shedding from Disp-deficient cells was restored by pharmacological membrane cholesterol extraction and by overexpressed transgenic Disp or structurally related Patched (Ptc, a putative cholesterol transporter). These data suggest that Disp regulates physiological Shh function via controlled cell surface shedding and that molecular mechanisms shared by Disp and Ptc exercise such sheddase control.

## Introduction

Hedgehog (Hh) ligands activate an evolutionarily conserved signaling pathway that provides instructional cues during tissue morphogenesis and, if misregulated, can contribute to developmental disorders and cancer. Several features of Hh signaling are highly unusual. The first feature is the covalent autocatalytic addition of a cholesteryl moiety to the C-terminal (Porter, Young, & Beachy, 1996) and of a palmitoyl group to the N-terminal by separate Hh acyltransferase (Hhat) activity (Pepinsky et al., 1998) (Fig. 1A). Hh N-palmitoylation maximizes Hh biofunction *in vivo* (Chamoun et al., 2001), and both lipids firmly tether Hh to the plasma membrane of the producing cell as a prerequisite for their multimerization (Vyas et al., 2008) and to effectively prevent unregulated ligand release. Signaling at distant cells therefore requires the removal of multimeric Hh from the membrane, a process that is facilitated by vertebrate and invertebrate Dispatched (Disp) orthologs: genetic studies in flies and mice revealed that Disp is specifically required in Hh ligand-producing cells and that Disp inactivation reduces ligand release and compromises Hh pathway activity *in vivo* (Burke et al., 1999; Kawakami et al., 2002; Ma et al., 2002; Nakano et al., 2004). Yet, the mechanistics of Disp-dependent Hh release remained unclear. Long-lasting questions about the Hh pathway are therefore i) how Disp drives dual-lipidated Hh release from the plasma membrane, ii) whether Disp acts directly or indirectly in the process, and iii) to what carrier – if any – Hh is transferred.

**Figure 1:**
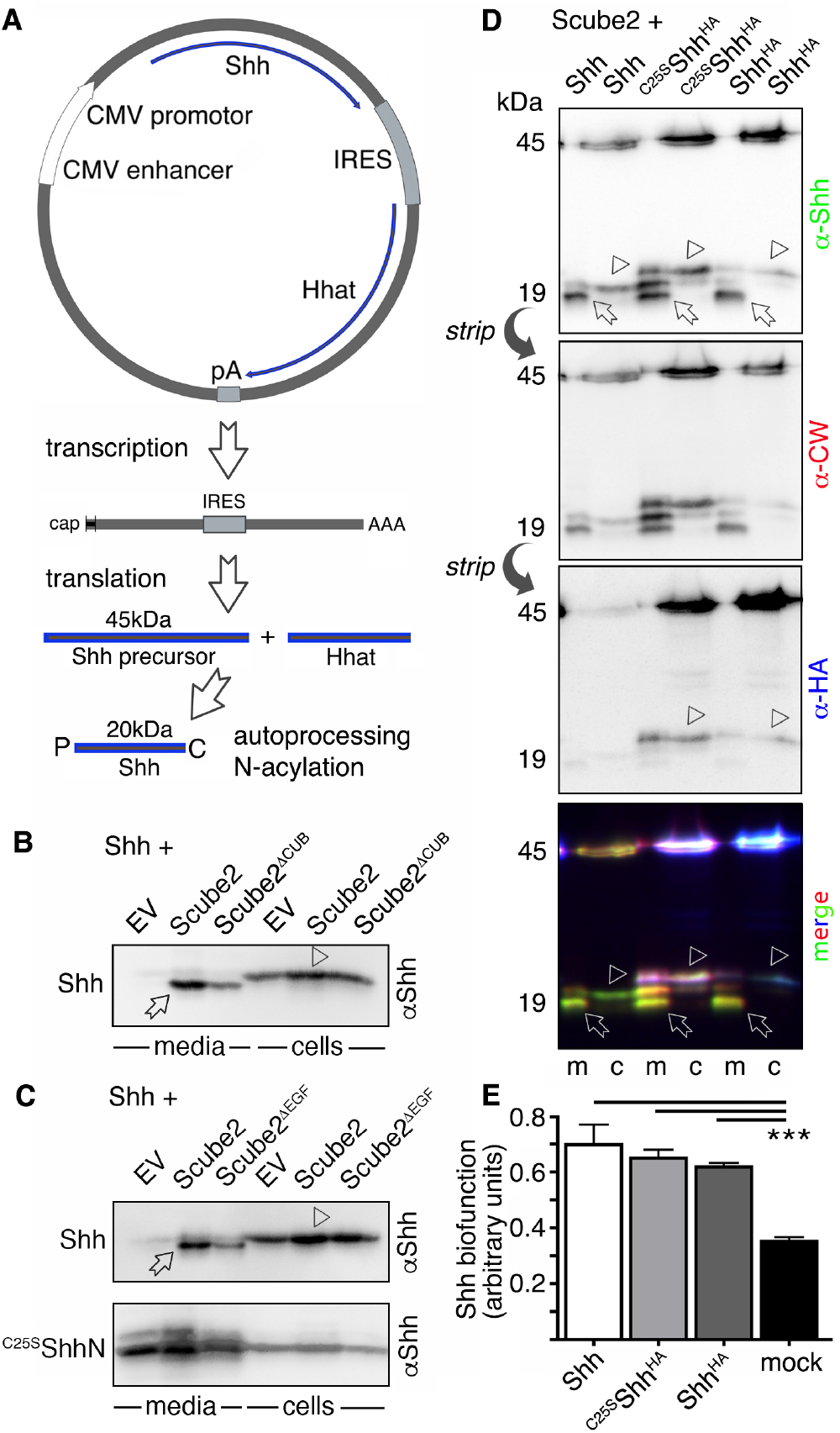
Dual-lipidated Shh release by Scube2-enhanced shedding. **A)** Cap-dependent Shh translation and cap-independent Hhat translation from bicistronic mRNA. IRES: internal ribosomal entry site. The 45 kDa Shh precursor autoprocesses into the cholesteroylated (C) 20 kDa Shh signaling domain. Shh N-palmitoylation (P) is independently catalyzed by Hhat. **B,C)** Scube2 enhances the conversion of cellular, dual-lipidated Shh (arrowhead) into truncated soluble forms (arrow). Scube2^ΔCUB^ and Scube2^ΔEGF^ are less active. Unlipidated ^C25S^ShhN is secreted independent of Scube2. EV: empty vector control. **D)** Co-expressed Scube2 increases conversion of tagged and untagged Shh variants (arrowheads) into similarly sized truncated proteins (arrows). Truncated Shh lacked α-HA reactivity and showed reduced antibody reactivity against an N-terminal peptide sequence called the Cardin-Weintraub (CW) motif. This indicates proteolytic cleavage upstream or within these sites, resulting in the loss of C-terminal and N-terminal peptides. Emergence of an upper band corresponding in size to the cellular signal (arrowheads) suggests a new processing site downstream of the HA tag (close to the cholesterol, see also Fig. S1). The same (stripped) blot was used for all incubations. Bottom: Merge of inverted, false-colored α-Shh, α-CW, and α-HA immunoblots. **E)** All truncated protein variants are bioactive. ***: p<0.001, n=2. One-way analysis of variance (ANOVA), Dunnett’s multiple comparison post hoc test.

What makes these questions particularly interesting is that the Hh release protein Disp on Hh producing cells is structurally related to the Hh receptor Patched (Ptc) on Hh receiving cells (Hall, Cleverdon, & Ogden, 2019). Both proteins contain 12 transmembrane helices and 2 extracellular domains and belong to the resistance-nodulation-division (RND) family of transmembrane efflux pumps. Normally, such pumps maintain cellular homeostasis and remove toxic compounds. In addition, both proteins contain a conserved domain known as the sterol-sensing domain (SSD) that is involved in different aspects of homeostasis of free or esterified cellular cholesterol in other SSD proteins (Hall et al., 2019). These striking structural resemblances between Ptc and Disp and the conserved SSD constitute another unexplained feature of the Hh pathway because they imply that similar – possibly cholesterol related – mechanisms control the opposite functions of Hh release from producing cells and Hh perception at receiving cells.

Notably, Disp activity alone is insufficient to deploy the vertebrate Hh family member Sonic hedgehog (Shh) from the plasma membrane of cells that express it. A second required synergistic factor required for maximum Shh signaling has been identified as the secreted glycoprotein Scube2 (Signal sequence, cubulin (CUB) domain, epidermal growth factor (EGF)-like protein 2) (Hall et al., 2019). One way to explain this synergy is that Disp may be required to extract lipidated Shh from the plasma membrane and hand it over to Scube2. The resulting stable Scube2-Shh complex may then relay and signal to receiving cells that express the receptor Ptc. This model has been supported *in vitro* by Disp- and Scube2 co-immunoprecipitation with Shh (Creanga et al., 2012; Tukachinsky, Kuzmickas, Jao, Liu, & Salic, 2012), and recently published Scube2-Shh interactions with Ptc coreceptors Cdon/Boc and Gas1 (Wierbowski et al., 2020). Notably, these activities critically depend on the C-terminal cysteine-rich and CUB domain of Scube2 (Creanga et al., 2012; Tukachinsky et al., 2012). CUB domains, however, derive their name from the complement subcomponents <u>C</u>1r/C1s, <u>s</u>ea urchin protein with EGF-like domains (UEGF), and <u>b</u>one morphogenetic protein 1 (BMP1) and contribute to protease activities in these proteins (Gaboriaud et al., 2011), possibly by inducing structural changes in the substrate to boost turnover (Bourhis et al., 2013; Jakobs et al., 2017). In agreement with this established CUB function, a suggested alternative role of Scube2 is to strongly enhance proteolytic removal (shedding) of both terminal lipidated Shh peptides to solubilize the morphogen in CUB-domain dependent manner (Jakobs et al., 2014; Jakobs et al., 2016; Jakobs et al., 2017). Shh shedding itself was suggested to require expression of A Disintegrin and Metalloproteinase family members 10 (A10) and 17 (Damhofer et al., 2015; Dierker, Dreier, Petersen, Bordych, & Grobe, 2009; Ohlig et al., 2011), two major sheddases known to cleave numerous substrate ectodomains from the cell surface. In this model, the requirement of Hh N-palmitoylation for maximum Hh signalling is only indirect: First, N-palmitate facilitates shedding by positioning the N-terminal peptide close to the cell surface (Baran, Nitz, Grotzinger, Scheller, & Garbers, 2013; Scheller, Chalaris, Garbers, & Rose-John, 2011; Zhao, Shey, Farnsworth, & Dailey, 2001). Second, with the continued membrane association of these N-terminal peptides, palmitate limits possible modes of morphogen solubilization to shedding and assures completion of this process. This is important, because unprocessed palmitoylated peptides sterically inhibit Shh binding sites for the receptor Ptc, both *in vitro* (Ohlig et al., 2011) and *in vivo* (Schürmann et al., 2018). Therefore, N-terminal proteolytic processing not only releases Shh but also unmasks Ptc binding sites and thereby couples Shh solubilization with its bioactivation. As a consequence, lack of acyltransferase activity in mutant cells or animals generates soluble inactive morphogens with (most) Ptc binding sites still blocked by their unprocessed N-termini. This hypothesis gained recent *in vivo* support: using *Drosophila melanogaster* wing- and eye development as models, site-directed mutagenesis of the predicted N-terminal sheddase cleavage site inactivated palmitoylated recombinant Hhs to variable degrees, depending on the mutation. Even more strikingly, sheddase-resistant transgene expression together with endogenous Hh in the same compartment resulted in strong dominant-negative wing phenotypes, probably caused by endogenous Hh trapping together with the sheddase-resistant transgenes in the same cell surface clusters (Kastl et al., 2018; Schürmann et al., 2018). *In vivo*, dominant-negative phenotypes were reversed by the additional removal of the N-palmitate membrane anchors from the transgenes, further supporting that proteolytic processing of N-terminal Hh membrane anchors controls Hh solubilization and biofunction. Finally, proteolytic processing of cholesteroylated C-termini has also been observed (Jakobs et al., 2014; Jakobs et al., 2016; Jakobs et al., 2017). We note that the C-terminal Hh target peptide is also likely cleaved during Hh release *in vivo*, because its replacement with transmembrane-CD2 (which is no known sheddase substrate) abolishes all Hh biofunction during *Drosophila* wing development (Strigini & Cohen, 1997).

In this study, we used CRISPR/Cas9 to determine the relation of Shh shedding to the established role of Disp in increasing Hh release. To this end, we compared the release of Shh from Disp-deficient Bosc23 cells with that of Disp-expressing control cells. We found that in the presence of Scube2, Disp specifically regulates proteolytic Shh shedding from its lipid membrane anchors, most likely by controlling spatial distribution of membrane cholesterol and Hh signaling component compartmentalization at the cell surface. This hypothesis is consistent with recent cryo-electron microscopy-derived data suggesting that structurally related Ptc removes cholesterol from the inner leaflet of the plasma membrane (Zhang et al., 2018) to suppress the 7-pass, G-protein-coupled receptor Smoothened (Smo). In line with such Ptc function and striking structural similarities between Ptc and Disp, Disp may regulate the Hh pathway via modulating Shh sheddase activity or access to Hh signaling components in a cholesterol-dependent manner. We support this idea by restored Shh shedding from Disp-deficient Bosc23 cells upon pharmacological cholesterol depletion or overexpression of transgenic Ptc. In addition to providing first insight into indirect physiological Shh sheddase control by Disp-regulated membrane lipid composition or distribution, our data link the known structural conservation between Disp and Ptc with a shared molecular mechanism that is essential for both: Hh relay from producing cells and Hh perception in target cells.

## Results

### Scube2 increases Shh shedding from Disp-expressing cells

To analyze whether Shh is released from the plasma membrane of producing cells via Scube2 CUB-controlled proteolysis or via Scube2 association, we produced Shh in Bosc23 cells, a derivative of HEK293 cells that endogenously express Disp (Jakobs et al., 2014). Shh biosynthesis in these cells requires several important posttranslational modifications to generate the dual-lipidated, plasma membrane-associated morphogen. The first modification consists of the removal of the Shh signal sequence during export into the endoplasmic reticulum. The resulting 45 kDa precursor proteins consist of an N-terminal signaling domain that starts with a cysteine (C25 in mouse Shh) and a C-terminal autoprocessing/cholesterol transferase domain. In a unique reaction, the autoprocessing/cholesterol transferase domain covalently attaches cholesterol to the C-terminus of the N-terminal signaling domain and simultaneously splits the 45 kDa precursor protein at the cholesteroylation site (Bumcrot, Takada, & McMahon, 1995). This coupled reaction ensures quantitative C-terminal cholesteroylation of all Shh signaling domains. In contrast, N-lipidation of signaling domains requires a separate enzymatic activity encoded by the Hh palmitoyltransferase Hhat, the lack or insufficient expression of which results in the secretion of non-palmitoylated Shh (Chamoun et al., 2001). Because HEK293 and Bosc23 cells do not express endogenous Hhat (Jakobs et al., 2014), throughout this work, we minimized the production of non-palmitoylated or only partially palmitoylated overexpressed Shh by using bicistronic mRNA to express the 45 kDa Shh precursor together with Hhat (Fig. 1A). Unlike Shh expression in its absence, Shh/Hhat co-expression from the same mRNA ensures near-quantitative Shh N-palmitoylation (Jakobs et al., 2014), which results in firm Shh plasma membrane association via both terminal peptides. We then analyzed how dual-lipidated, membrane-associated Shh is released from the plasma membrane into the supernatant by SDS-PAGE/immunoblotting. As shown in Figure 1B, Scube2 enhanced Shh release from Bosc23 cells, as expected. Notably, we also observed that the electrophoretic mobility of most released Shh was increased over that of the corresponding dual-lipidated cellular material (Jakobs et al., 2014; Jakobs et al., 2016). Finally, and as previously noted (Jakobs et al., 2014; Jakobs et al., 2017), we confirmed that Scube2 variants lacking their CUB or EGF domains release less truncated Shh and that engineered soluble Shh lacking both lipids (^C25S^ShhN) was always released in unprocessed form and independent of Scube2 (Fig. 1B,C). The observed increase in electrophoretic mobility of most soluble Shh over its lipidated cellular precursor can be best explained by the proteolytic removal of both lipids together with the associated terminal peptides (shedding; details are outlined in Fig. S1). We supported Scube2-enhanced Shh shedding by the analysis of hemagglutinin (HA)-tagged palmitoylated Shh^HA^ and non-palmitoylated ^C25S^Shh^HA^ variant proteins. In both proteins, the HA tag was inserted adjacent to the cholesteroylated glycine 198 (Fig. 1D). We observed that palmitoylated cellular Shh and Shh^HA^ converted into similar truncated proteins upon solubilization. Yet, N-terminal processing of non-palmitoylated ^C25S^Shh^HA^ was markedly reduced, as indicated by the appearance of additional soluble products with decreased electrophoretic mobility (labelled orange and violet in the merged, false-colored blot). Their appearance suggests that palmitate, via its continued membrane association, controls quantitative N-terminal Shh processing during release and that the minor amount of N-terminally unprocessed soluble proteins therefore represent the Shh fraction that did not undergo N-palmitoylation during biosynthesis (Fig. S1). This finding underlines the importance of Hhat co-expression to properly characterize Shh release. Notably, if normalized for the fully processed protein amounts (the bottom media bands, labelled green in the merged blot, arrows), all soluble constructs showed similar bioactivities as determined by differentiation of the Shh-responsive cell line C3H10T1/2 into alkaline phosphatase-producing osteoblasts (Ohlig et al., 2011). Because ^C25S^Shh^HA^ was never palmitoylated, this observation confirms that lipidation-dependent Shh processing, and not Shh palmitoylation *per se*, determines Scube2-regulated Shh signaling activities in receiving cells (Ohlig et al., 2011) (Fig. 1E).

### Disp is specifically required for Scube2-regulated Shh shedding

Because Scube2-regulated Hh release is known to also require Disp (Hall et al., 2019), we expected strongly impaired Shh shedding from cells made deficient in Disp function. To test this hypothesis, we generated Disp knockout cells (Disp^−/−^) by using CRISPR/Cas9. Sequencing of the targeted genomic loci confirmed deletion of 7 base pairs, leading to a frameshift and a stop codon at amino acid 323 located in the first extracellular loop of the 1524 amino acid protein (Fig. 2A, A’). We verified unaffected predicted off-targets (Table S1) and confirmed complete Disp protein loss in Disp^−/−^ cells via immunoblotting (Fig. S2A). Transgenic Disp overexpression (Disp^tg^) restored the immunoblot signal, demonstrating effectivity and specificity of the targeting approach (Fig. S2A, arrows). Consistent with our expectations, Shh shedding from Disp^−/−^ cells was significantly reduced compared with truncated Shh release from Shh-expressing *CRISPR non*-*targeting* control cells (nt Ctrl) (Figure 2B arrows, B’). Instead, Disp^−/−^ cells accumulated cellular Shh (Fig. 2B, arrowhead), similar to what has been previously shown *in vivo* (Burke et al., 1999) (Fig. 2B, arrowhead). Also consistent with previous observations (Kawakami et al., 2002; Ma et al., 2002), loss of Disp in Bosc23 cells did not affect Shh biosynthesis, autoprocessing, and coupled cholesteroylation of the 19 kDa N-terminal signaling domain (Fig. S2B), ruling out any role of Disp in these processes. Disp loss also did not affect morphogen secretion, because unlipidated control ^C25S^ShhN was readily released (Fig. 2C arrows, C’). We conclude that Disp specifically increases shedding of bioactive Shh (Fig. S3) from the surface of producing cells, but that Disp is not essential for this process *per se*, because small amounts of processed Shh are still released from Disp^−/−^ cells. Required yet non-essential Disp regulation of Shh shedding supports published observations that Disp is not absolutely required for paracrine Indian Hh (a Shh homolog) signaling in the skeleton (Tsiairis & McMahon, 2008), that overexpression of Shh can overcome Disp restriction (Nakano et al., 2004), and that cells or tissues that generate high levels of Hh are partially independent of Disp (Burke et al., 1999; Caspary et al., 2002; Ma et al., 2002). Of note, these reports of required but non-essential Disp function in Shh release do not support the possibility of direct Disp binding of Hh cholesterol to hand the lipidated (unprocessed) Shh over to Scube2, as Disp deletion should completely block this process. Moreover, in the absence of Scube2 (Fig. 2D, D’) or in the presence of serum (Fig. 2E, E’), apparently unprocessed Shh (represented by the “upper” band, arrows) is somehow released in a Disp-independent manner, possibly still associated with membranous carriers. Disp-independent release of unprocessed Shh is therefore not likely of physiological relevance, and such proteins have indeed been described as inactive before (Palm et al., 2013; Tukachinsky et al., 2012). In contrast, Scube2 and Disp synergistically increase Shh shedding, which strongly suggests a physiological role of truncated soluble Shh release.

**Figure 2:**
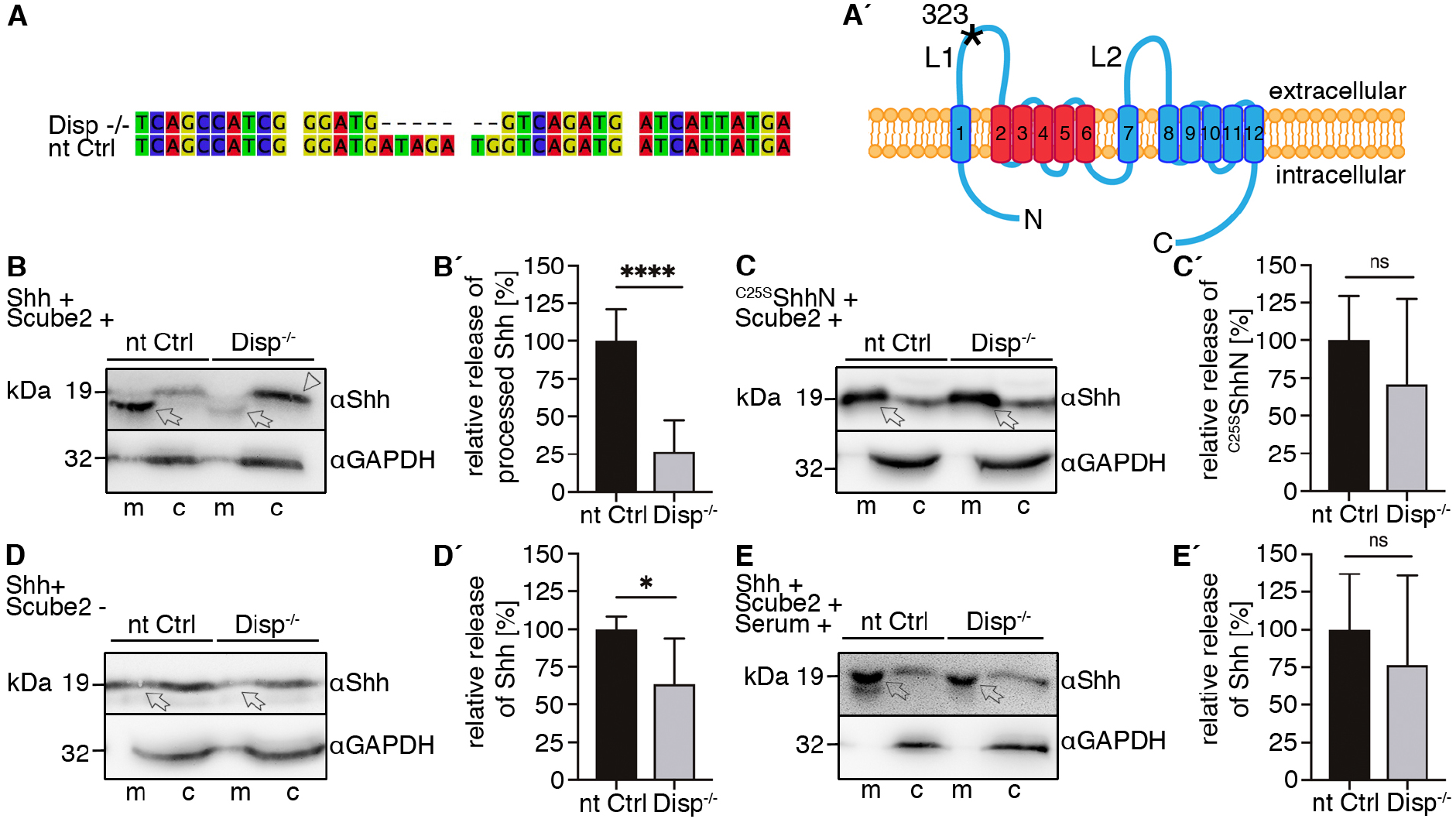
Impaired but not abolished Shh release from Disp^−/−^ cells. **A)** Alignment of targeted *disp* gene sequences from Disp^−/−^ cells and from non-targeted (nt Ctrl) cells. **A’)** Schematic representation of the Disp protein structure. An asterisk indicates the CRISPR/Cas9-generated stop codon introduced at position 323. This deleted 11 of 12 transmembrane domains (TM), representing ~80% of the protein sequence. L1/L2: extracellular loops. TM2-6 (red): sterol-sensing domain (SSD). **B-E)** Immunoblots of cellular (c) and released (into the media, m) Shh and unlipidated control ^C25S^ShhN in nt Ctrl and Disp^−/−^ cells. Scube2 co-expression is indicated; arrows indicate solubilized Shh and the arrowhead indicates accumulated cellular material in Disp^−/−^ cells. D, E: The absence of Scube2 in serum-free media or the presence of 10% serum reduced Shh processing and rendered Shh release independent of Disp. **B’-E’)** Quantifications of relative Shh release from nt Ctrl and Disp^−/−^ cells. Ratios of solubilized versus cellular Shh were determined and expressed relative to Shh release from nt Ctrl cells (set to 100%, black bars). Unpaired t-tests. B’: n=22 datasets from 11 independent experiments. C’: n=9 datasets from 4 independent experiments. D’: n=7 datasets from 3 independent experiments. E’: n=18 datasets from 5 independent experiments.

We next investigated the relative roles of N-palmitate and C-cholesterol in Disp-regulated Shh shedding and found that the release of palmitoylated/non-cholesteroylated ShhN and of cholesteroylated/non-palmitoylated ^C25A^Shh did not strictly depend on Disp (Fig. 3A-B’). Again, this finding supports the importance of dual Hh lipidation for its proper *in vitro* characterization and provides an explanation for the requirement for, and complete conservation of, N- and C-terminal Hh lipidation in vertebrates and invertebrates. Finally, to confirm that impaired Shh shedding was specifically caused by disrupted Disp function and not through any off-target events that might have been missed in our confirmatory screen (Table S1), we aimed to reverse the Disp^−/−^ phenotype by the expression of transgenic V5-tagged Disp^tg^ in these cells. Confocal microscopy of non-permeabilized Disp^−/−^ cells expressing either Shh or Disp^tg^ confirmed secretion of both proteins to the cell surface (Fig. 3C, C’). Consistent with their cell-surface localization, co-expressed Disp^tg^ restored Shh shedding from Disp^−/−^ cells (Fig. 3D arrows, D’). On the basis of the recently resolved cryo-electron microscopy structure of *Drosophila* Disp (Cannac et al., 2020), we also tested the activity of a Disp^ΔL2^ variant lacking amino acids 752-972 of the second extracellular loop, located between transmembrane (TM) regions 7 and 8 (Fig. 2A’). In the cryo-electron microscopy structure, both Disp extracellular loops adopted an open conformation, and 3-dimensional reconstruction allowed Hh ligand binding to both loops (Cannac et al., 2020). We indeed observed reduced Disp^ΔL2^-mediated Shh shedding from Disp^−/−^ cells if compared to full-length Disp (Fig. 3D, D’), suggesting a non-essential role of the second extracellular loop in Shh release. Shh shedding from Disp^tg^ and DispΔL2 transfected nt Ctrl cells was not significantly increased, supporting sufficient endogenous Disp expression in these cells (Fig. 3E,E’). From these observations, we conclude that Disp releases Shh by controlling Shh shedding from the plasma membrane in a DispL2-dependent manner.

**Figure 3:**
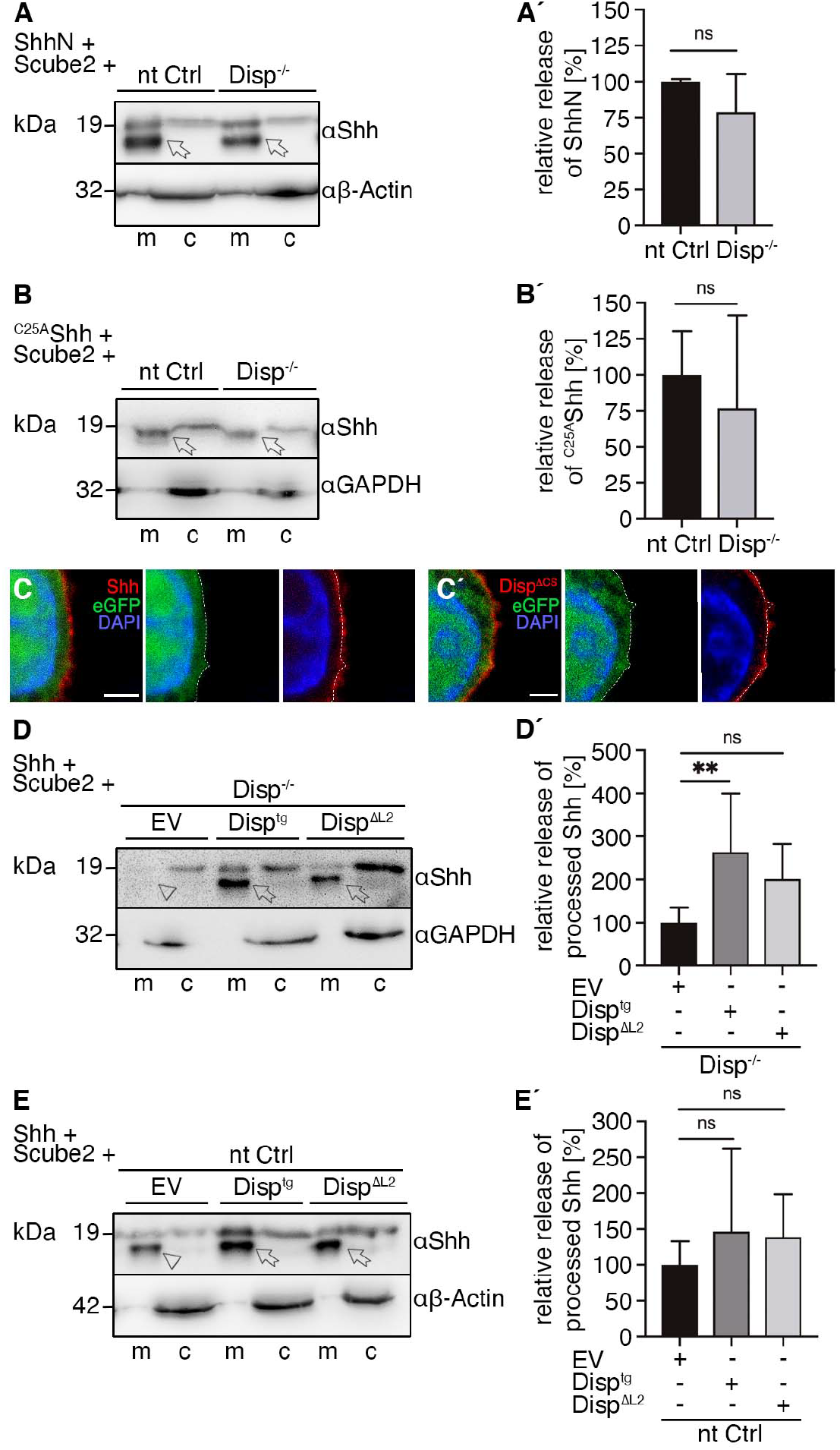
Disp^tg^ expression rescues Shh release from Disp^−/−^ cells. **A, A’)** Similar relative release of N-palmitoylated but non-cholesteroylated ShhN into serum-free media (m) from nt Ctrl and Disp^−/−^ cells (c). Arrows indicate solubilized processed Shh. n=5 datasets from 4 independent experiments, unpaired t-test. **B, B’)** Similar relative release of non-palmitoylated but cholesterol-modified ^C25A^Shh from nt Ctrl and Disp^−/−^ cells. Arrows indicate solubilized Shh. n=11 datasets from 3 independent experiments, unpaired t-test. **C, C’)** Representative confocal planes of Disp^−/−^ cells expressing either Shh (C, red) or Disp^tg^ (C’, red). Nuclei are counterstained with DAPI and eGFP served to visualize the cytoplasm. Dashed lines indicate the border of cytoplasmic GFP signals. Both transgenes were secreted to the cell surface. Scale bars: 2 μm. **D, D’)** Co-expressed transgenic Disp^tg^ enhanced processed Shh release from Disp^−/−^ cells, and transgenic Disp lacking most of the second extracellular loop (Disp^ΔL2^) did not significantly increase Shh release. D’: Quantified relative Shh release as shown in D. n=9 datasets from 5 independent experiments, one-way ANOVA, Dunnett’s multiple comparison post hoc test. **E, E’)** Co-expressed transgenic Disp^tg^ or Disp^ΔL2^ did not significantly increase Shh release from nt Ctrl cells. E’: Quantified relative Shh release as shown in E. n=12 datasets from 6 independent experiments, one-way ANOVA, Dunnett’s multiple comparison post hoc test. D, E: Arrows indicate solubilized Shh from Disp^tg^-or Disp^ΔL2^-expressing cells and the arrowhead indicates solubilized Shh from empty vector (EV) transfected cells.

### Loss of Disp function is restored by expression of the putative cholesterol transporter Ptc

How can Shh shedding control by Disp be explained? One solution to this question comes from established structural similarities between Disp and Ptc: both are 12-TM proteins that contain a sterol-sensing domain (SSD, consisting of TM2-TM6, Fig. 4A, A’, colored red) and 2 extracellular loops. More recently, cryo-electron microscopy revealed that the SSDs of Disp and of Ptc can be superimposed and accommodate several sterol-like densities in a central hydrophobic conduit (Cannac et al., 2020; Gong et al., 2018). The Ptc SSD also possesses similarities to prokaryotic RND transporters that function as proton-driven antiporters, often to export molecules through a hydrophobic channel that resembles the conduit of Ptc (Nies, 1995; Tseng et al., 1999). This finding indicated that Ptc may use this conduit to transport cholesterol, and indeed, active Ptc was shown to decrease free cholesterol amounts (Bidet et al., 2011), in particular in the inner plasma membrane leaflet to keep Smo in an inactivate state (Zhang et al., 2018). Moreover, as suggested for Disp (Cannac et al., 2020), L1 and L2 of Ptc constitute the Hh binding site, and Hh association with this site stops Ptc-mediated cholesterol transport. This, in turn, leads to inner-leaflet cholesterol accumulation and Smo activation (Zhang et al., 2018). From these findings, we hypothesized that Disp may likewise transport free (unesterified) cellular cholesterol, 60%-80% of which resides in the plasma membrane. We also hypothesized that, if that is correct, Ptc^tg^ co-expression, together with Shh/Hhat and Scube2, may compensate for Disp loss in our mutant cell line. To test these hypotheses, we co-transfected Disp^−/−^ cells and nt Ctrl cells with Ptc^tg^ or constitutively active Ptc^ΔL2^ that lacks Shh-binding L2, which renders the molecule insensitive to activity down-regulation by Shh (Taipale, Cooper, Maiti, & Beachy, 2002). As expected, Ptc^tg^ and Ptc^ΔL2^ strongly increased Shh shedding from Disp^−/−^ cells (Fig. 4B arrows, B’), as well as from control cells (Fig. 4C, 4C’). This observation suggests that Ptc and Disp act in similar mechanistic manner, consistent with their structural conservation (Cannac et al., 2020).

**Figure 4:**
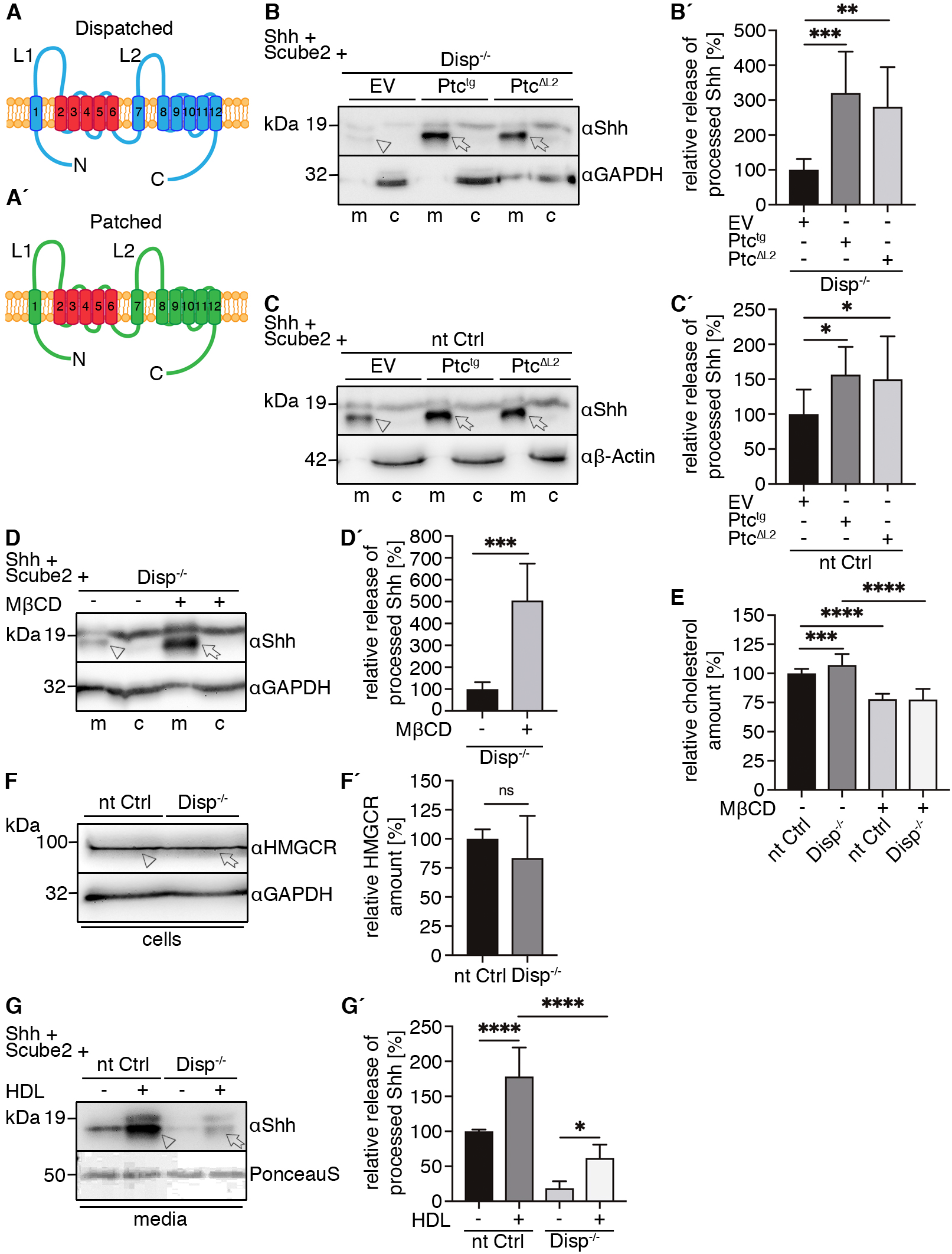
Cellular cholesterol depletion induces Shh shedding. **A, A’)** Schematic representation of Disp (blue) and Ptc (green). Twelve transmembrane domains (TM1-12), 2 extracellular loops (L1 and L2), and the N- and C-termini are labelled. The sterol-sensing domains (SSDs, TM2-6) are highlighted in red. **B, C)** Co-expression of transgenic Ptc^tg^ or a Ptc variant lacking most of the second extracellular loop (Ptc^ΔL2^) increases Shh shedding from Disp^−/−^ (B) and nt Ctrl (C) cells. Arrows indicate solubilized processed Shh from Ptc^tg^-or Ptc^ΔL2^-expressing cells and the arrowhead indicates solubilized Shh from empty vector (EV) transfected cells. **B’, C’)** Quantification of relative processed Shh release as shown in B and C. B’: n=10 datasets from 5 independent experiments. C’: n=11 datasets from 6 independent experiments, one-way ANOVA, Dunnett’s multiple comparison post hoc test. **D, D’)** Shh shedding is enhanced by the cholesterol-depleting drug MβCD. Solubilized Shh in the absence or presence of MβCD is indicated by an arrowhead and an arrow, respectively. n=6 datasets from 3 independent experiments, unpaired t-test. **E)** Quantified relative free cholesterol content in nt Ctrl and Disp^−/−^ cells in the presence or absence of MβCD. n=29 datasets from 5 independent experiments, one-way ANOVA, Sidak’s multiple comparison post hoc test. **F, F’)** Similar HMGCR expression in nt Ctrl cells (arrowhead) and Disp^−/−^ cells (arrow). n=8 datasets from 3 independent experiments, unpaired t-test. **G)** The presence of HDL enhanced Shh release from nt Ctrl cells (arrowhead) but less so from Disp^−/−^ cells (arrow). Ponceau S staining served as a media loading control. **G’)** Quantification of relative processed Shh release. n=6 datasets from 3 independent experiments, one-way ANOVA, Sidak’s multiple comparison post hoc test.

To test whether this shared mechanism is related to the postulated cholesterol transporter-like activity of Ptc, we depleted Disp^−/−^ cells of free membrane cholesterol by using the cholesterol-depleting drug *methyl*-*β*-*cyclodextrin* (MβCD) (Zidovetzki & Levitan, 2007). Indeed, MβCD restored Shh shedding from Disp^−/−^ cells in a concentration-dependent manner (Fig. 4D arrow, D’, Fig. S4), showing that pharmacologically decreased membrane cholesterol increases Shh shedding and suggesting that Ptc and Disp may regulate free membrane cholesterol content in a physiologically relevant manner. To further investigate this possibility, we quantified total (esterified) and free cholesterol in Disp^−/−^ and nt Ctrl cells. In line with its established function as a membrane cholesterol extractor (Zidovetzki & Levitan, 2007), MβCD significantly reduced both cholesterol pools in our assay (Fig. 4E, Fig. S5). We hypothesized that, if Disp acted in the same manner, we should observe an increase in membrane cholesterol in Disp^−/−^ cells compared with nt Ctrl cells. Our quantitative assay supported this hypothesis because free cholesterol amounts in Disp^−/−^ cells were significantly, yet only slightly, increased over those in nt Ctrl cells (Fig. 4E, Fig. S5). We also analyzed a key enzyme of *de novo* cholesterol biosynthesis called 3-hydroxy-3-methyl-glutaryl coenzyme A reductase (HMGCR). HMGCR expression can serve as a read-out for cellular cholesterol content because sterol depletion in the endoplasmic reticulum stimulates HMGCR synthesis, and high cholesterol levels induce rapid HMGCR degradation (Kuwabara & Labouesse, 2002; Luo, Yang, & Song, 2020). Similar HMGCR in Disp^−/−^ and nt Ctrl cells showed that the observed slight increase in cholesterol did not affect its *de novo* biosynthesis (Fig. 4F, F’). We also used qPCR to investigate all aspects of cholesterol homeostasis: here, we again detected only slight reductions in the transcription of genes regulating *de novo* cholesterol biosynthesis, dietary cholesterol uptake via Niemann-Pick C1-like protein 1, low-density lipoprotein (LDL)-mediated cholesterol uptake, cholesterol efflux, and cholesterol storage via esterification (Fig S6). The only exception was upregulated proprotein convertase subtilisin/kexin type 9 (PCSK9) transcription in Disp^−/−^ cells. Because one role of PCSK9 is to degrade the LDL receptor, and LDL receptor degradation reduces cellular cholesterol uptake, the observed increase in PCSK9 transcription may indicate a response to the slight increase in cellular cholesterol. Taken together, our findings suggest that Disp may regulate Shh shedding by changing membrane cholesterol, probably at sites of Shh storage at the cell surface. Such local Disp function would again resemble local Ptc activities, which are restricted to the membrane environment of the primary cilium in vertebrates.

### Disp function requires high-density lipoproteins

At the primary cilium, Ptc depletes free cholesterol from the inner plasma membrane leaflet, yet the site of cholesterol deposition has remained unidentified (Zhang et al., 2018). One possibility is that the cholesterol is merely transferred to the outer membrane leaflet of the plasma membrane. In this case, Ptc would act as a cholesterol floppase. The alternative possibility is cholesterol export to a sink – a soluble carrier to which the sterol is transferred and transported away from the cell. In invertebrates, a known cholesterol sink is lipoproteins called lipophorins. In vertebrates, high-density lipoproteins (HDLs) are a specialized lipoprotein fraction that accepts peripheral cholesterol from transmembrane transporters of the ABC family for its further use in steroid hormone-producing cells or for secretion by the liver (Luo et al., 2020). This function would make HDL suited to accepting the sterol-like densities present in the conserved SSDs of Ptc and Disp (Cannac et al., 2020; Zhang et al., 2018). To test this potential, and to strengthen the suspected link between physiological cholesterol export and Shh shedding, we added purified human HDL to serum-free media of Shh-expressing Disp^−/−^ cells and nt Ctrl cells. We expected that added HDL would increase Shh shedding from nt Ctrl cells and that Disp^−/−^ cells would be less responsive due to the lack of the Shh-specific Disp exporter. Indeed, HDL significantly increased Shh shedding from nt Ctrl cells (Fig. 4G arrowhead, G’, Fig. S7) and to a much lesser degree from Disp^−/−^ cells (Fig. 4G arrow, G’, Fig. S7). Weakly enhanced Shh shedding from Disp^−/−^ cells was not unexpected due to the expression of ABC cholesterol exporters to maintain cellular cholesterol homeostasis (Table S2): these also use HDL or discoidal precursor HDL as cholesterol acceptors (Luo et al., 2020) and thus may have partially compensated for Disp loss in our setting. Of note, the alternative possibility of direct cellular Shh extraction from the membrane to HDL acceptors is not supported, as this should have left electrophoretic mobility of soluble Shh unchanged. Instead, our data suggest that established extracellular cholesterol carriers such as HDL may act downstream of Disp to accept free membrane sterol. This, in turn, may decrease local cholesterol concentration or distribution at the cell surface to stimulate Shh shedding.

### Shh sheddase activity is regulated by its transmembrane domain

One hypothesis to explain how locally changed cholesterol affects shedding is via regulation of sheddase access to its substrates. It is known that artificial membrane cholesterol depletion by MβCD can non-specifically activate cell-surface shedding (Reiss & Bhakdi, 2017), and one established MβCD-responsive, membrane cholesterol-dependent sheddase is A10 (Kojro, Gimpl, Lammich, Marz, & Fahrenholz, 2001; Matthews et al., 2003; Murai et al., 2011). We confirmed (Ohlig et al., 2011) that cells deficient in A10 expression released less Shh than control cells did (Fig. 5A, arrows, A’). This was again in notable contrast to non-lipidated ^C25S^ShhN, the release of which was independent of A10 (Fig. 5B, arrows). We further supported the postulated link between Disp function and Shh shedding by Disp^tg^ expression in A10^−/−^ cells: This indeed restored Shh processing and release (Fig. 5C arrow, C’) when compared with A10^−/−^ cells not transfected with Disp^tg^ (Fig. 5C, arrowhead, C’). We explain A10 activity compensation by other cell-surface sheddases under the activating conditions of Disp-mediated membrane cholesterol depletion, as indicated by increased, yet different Shh processing in A10^−/−^ cells when compared with HEK control cells (WT, Fig. 5C, asterisk, C’).

**Figure 5:**
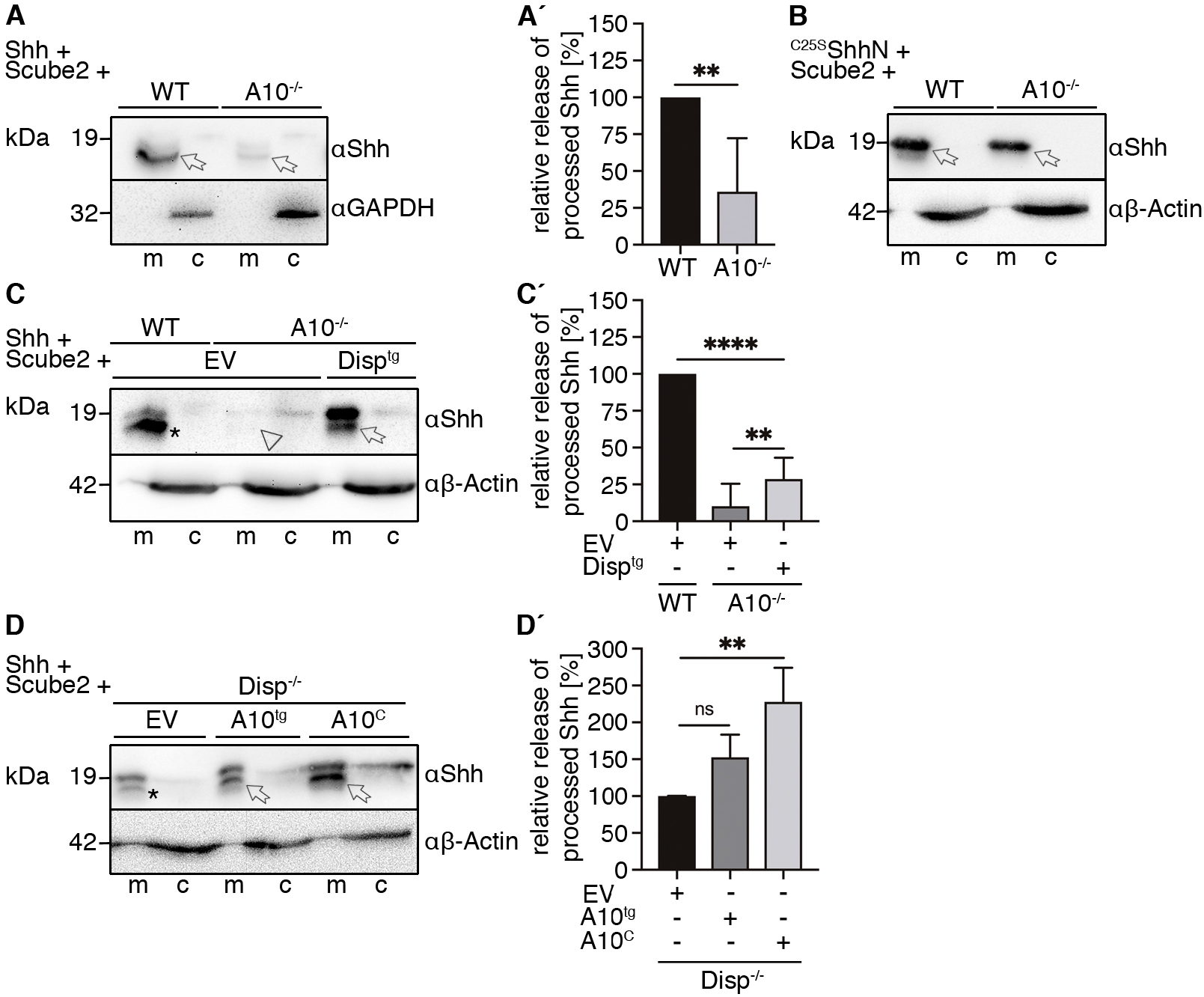
Loss of A10 function impairs Shh processing. **A)** Reduced Shh shedding from A10^−/−^ cells compared with Shh release from HEK wild-type (WT) cells. Arrows indicate solubilized truncated Shh. **A’)** Quantification of relative amounts of processed Shh as shown in A. n=5 datasets from 5 independent experiments, unpaired t-test. **B)** Similar unlipidated ^C25S^ShhN release from WT and A10^−/−^ cells (arrows). **C)** Increased release of processed Shh from A10^−/−^ cells that co-express Disp^tg^. Note the reduced release and only partial processing of Shh from A10^−/−^ cells (arrow and arrowhead) compared with empty vector (EV) transfected WT cells (asterisk). **C’)** Quantification of relative processed Shh release as shown in C. n=9 datasets from 4 independent experiments, one-way ANOVA, Sidak’s multiple comparison post hoc test. **D)** A10^C^ with its C-terminal transmembrane domain replaced by C-cholesterol increases Shh shedding from the surface of Disp^−/−^ cells (arrows). **D’)** Quantification of relative processed Shh release as shown in D. n=3 datasets from 3 independent experiments, one-way ANOVA, Dunnett’s multiple comparison post hoc test.

One attractive mechanism to explain the functional interdependency between membrane cholesterol and shedding is the postulated control of cell-surface signaling by cholesterol-enriched plasma membrane microdomains, called lipid rafts (Simons & Ikonen, 1997). Lipid rafts are plasma membrane substructures with a suggested average diameter of 100 nm to 200 nm that represent 13%-50% of the cellular surface, depending on the cell type. Rafts contain increased concentrations of cholesterol, sphingomyelin, and gangliosides, as well as phospholipids with saturated fatty acyl chains (Pike, 2003) that render them more densely packed and thus less fluid than the surrounding plasma membrane. From these biophysical properties, rafts are thought to control signaling by protecting incomplete signaling composites partitioned into rafts from activator components resident outside of rafts. Interestingly, extracellular proteins firmly established to partition into rafts are glycosylphosphatidylinositol-anchored proteins, including the Hh-associated heparan sulfate proteoglycans of the glypican family and dual-lipidated Hhs. Both molecules are special in that they insert only into the outer leaflet of the plasma membrane (Pike, 2003; Rietveld, Neutz, Simons, & Eaton, 1999). In contrast, A10 is a transmembrane protein and is thought to normally reside outside of rafts (Reiss & Bhakdi, 2017). Notably, such physical separation between raft and non-raft proteins at the cell surface becomes diminished on disintegration of the lipid raft, for example, as a consequence of MβCD treatment (Kojro et al., 2001; Simons & Ehehalt, 2002). The consequence of such a perturbed physical separation of raft ligands and non-raft release factors may have represented what we observed in our study, as MβCD-induced cholesterol depletion from the plasma membrane (Fig. 4E) strongly increased Shh release (Fig. 4D, D’). To directly test the hypothesis that rafts compartmentalize Hh/glypican assemblies (Vyas et al., 2008) to physically separate them from A10, we generated an A10^C^ protein variant with its transmembrane domain replaced by the ShhC autoprocessing/cholesterol transferase domain. Subsequent A10^C^ cholesteroylation during biosynthesis was expected to partition A10^C^ ectodomains into Shh-containing rafts, which would result in increased Shh shedding from Disp^−/−^ cells. In contrast, the TM domain of A10^tg^ may prevent access to rafts, which would render the protease less active due to its physical separation from the substrate. Increased Shh processing as a consequence of A10^C^ co-expression (Fig. 5D, arrow, D’), but less relative processing as a consequence of A10^tg^ co-expression (Fig. 5D, arrow, D’), supported our idea that rafts may provide a physical barrier between Hh ligands within rafts and the sheddase outside of rafts, and that increased raft size or half-life in Disp^−/−^ cells may boost and prolong this separation. We thus suggest that, in Hh signaling, rafts may be essential in limiting signaling by physical separation of Hh signaling components on producing cells until regulated Hh release is initiated.

## Discussion

Although Disp is firmly established as a critical component of the Hh pathway, 2 outstanding questions remained unanswered so far: i) What is the molecular mechanism by which Disp releases lipid-modified Shh from producing cell membranes? and ii) are there additional Disp functional partners to promote Shh deployment? As shown in this study, we provide answers to these questions and suggest a Hh release model that matches several predictions derived from previous genetic studies (Burke et al., 1999; Kawakami et al., 2002; Ma et al., 2002). Our model of Disp-modulated Shh shedding meets the first prediction that Disp is only required in Hh-producing cells and not in receiving cells. It meets the second prediction that Hh signaling defects in Disp-deficient model organisms are not due to a defect in Hh production or cholesteroylation, but rather to the deployment of lipidated (but not artificially produced unlipidated forms of) Shh from producing cells. Our data also support the third prediction that full Disp activity requires Scube2 (Tukachinsky et al., 2012) and can further explain why the Disp loss-of-function phenotype in *Drosophila* is less severe than that of the single Hh ortholog (Burke et al., 1999): we suggest that, while Disp is required for regulated Hh shedding, it is not absolutely essential, as has previously been shown for some Hh-producing tissue types *in vivo* (Caspary et al., 2002; Nakano et al., 2004; Tsiairis & McMahon, 2008) and as we now show for transfected cells that overexpress Shh *in vitro*. Finally, we demonstrate that established lipid carriers such as HDL can act as shedding enhancers, at least in our *in vitro* system. This finding suggests that Disp may extract cholesterol, or that it may contribute to a heteroprotein membrane complex that extracts the sterol from the plasma membrane as a prerequisite for its removal from the cell. In this regard, Disp might act like bacterial RND transporters and related Niemann Pick Type C protein 1 (an SSD transmembrane protein that transmits cholesterol to intracellular protein acceptors). If correct, our model, which suggests that Disp can export cholesterol and that lipoproteins can act as cholesterol sinks, would provide an explanation for the published observations that knockdown of fly lipophorin impairs Hh biofunction *in vivo* (Panakova, Sprong, Marois, Thiele, & Eaton, 2005), that cytonemes that transport Hh to target cells contain Disp at their tips (reviewed in (Hall et al., 2019)), and that Ptc acts as a lipoprotein receptor *in vivo* (Callejo, Culi, & Guerrero, 2008). Although our study did not directly address lipoprotein function in the fly, the possibility exists that Disp-mediated cholesterol relay to these carriers supports Hh shedding and signaling at cellular interfaces. This suggestion indicates that cholesterol not only stands out as one endogenous ligand of Ptc to regulate Smo in receiving cells (Zhang et al., 2018), but that it can also act as a physiological second messenger in Shh-producing cells. This possibility is supported by structural similarities between Ptc and Disp, including the presence of SSDs in both proteins.

We are aware that our work contrasts strongly with widely accepted models of direct contribution of Hh lipids to Ptc-mediated signaling. These models gained recent support by several cryo-electron microscopy-derived structures showing direct Hh lipid association with the Ptc-receptor. However, we point out that these structures were derived from conditions that always represented the Shh pre-release state, such as artificial co-expression of ligand and receptor in the same cells (Qian et al., 2019) or the reconstitution of detergent-solubilized Hh or Hh peptides with Ptc *in vitro* (Qi, Schmiege, Coutavas, & Li, 2018; Qi, Schmiege, Coutavas, Wang, & Li, 2018). The physiological relevance of such structures, therefore, depends on the question whether Disp and Scube2 - that both act upstream of Ptc - release Shh in unprocessed or in delipidated processed form. Support for the former possibility comes from three previous Shh overexpression studies (Creanga et al., 2012; Tukachinsky et al., 2012; Wierbowski et al., 2020), while this and previous work of our group demonstrate Shh release in delipidated processed form (Dierker et al., 2009; Jakobs et al., 2014; Jakobs et al., 2017; Ohlig et al., 2011; Ohlig et al., 2012; Schürmann et al., 2018). We explain these different findings by our Shh/Hhat co-expression system to generate physiologically relevant, dual-membrane associated Shh in near-quantitative manner. Only such proteins undergo proteolytic processing at both terminal peptides to generate truncated Shh, while N-terminally unmodified proteins that result from insufficient endogenous Hhat activity in Shh expressing cells do not (Figure S1)(Jakobs et al., 2014). As described before, this is because sheddases require membrane-proximal positioning of substrate cleavage sites for their activity, which is usually achieved by their membrane-association (Lichtenthaler, Lemberg, & Fluhrer, 2018). We also note that the current concept of lipidated Hh hand-over from Disp to Scube2 to GAS1 to Ptc (Creanga et al., 2012; Tukachinsky et al., 2012; Wierbowski et al., 2020) is in conflict with the *in vivo* observation that murine Disp expression in Disp-deficient *Drosophila* restores Hh dependent development (Burke et al., 1999), despite the lack of Scube orthologs in flies. It is also difficult to reconcile with the *in vivo* observation that transgenes with a 27kDa GFP tag inserted between the 19kDa Hh and its C-terminal cholesterol restore the development of Hh-deficient flies to adulthood (Chen, Huang, Hatori, & Kornberg, 2017), because this would require rather flexible mechanisms of Hh membrane extraction, repeated hand-over and receptor activation. Furthermore, Scube2 is not strictly required for Hh signaling in vertebrates, because the Scube2 mutant phenotype (called *you* phenotype in zebrafish) is the weakest in its class of mutants that disrupt the Hh signal transduction pathway, and because Shh overexpression can bypass Hh patterning defects as a consequence of Scube1-3 depletion *in vivo* (Johnson et al., 2012).These *in vivo* observations are incompatible with the concept of Scube2-mediated Shh transport, because any Scube1-3 transport blockade would not be bypassed by increased amounts of ligand. The concept of lipidated Hh hand-over also conflicts with several findings indicating that Hh can be released independent of Disp function (Burke et al., 1999; Caspary et al., 2002; Ma et al., 2002; Nakano et al., 2004; Tsiairis & McMahon, 2008), as well as with our finding that Disp function in dual-lipidated Shh release can be fully substituted by the receptor Ptc. Yet, every one of the above findings is compatible with Scube2- and Disp-modulated, sheddase-mediated Shh release off its lipidated terminal peptides, as described in our submitted study.

We also note that a second perceived problem with our shedding model is that the N-palmitate is known to increase Hh biofunction *in vitro* (Pepinsky et al., 1998) and *in vivo* (Chamoun et al., 2001; Kohtz et al., 2001; Lee et al., 2001), albeit to highly variable degrees. Current models explain this essential function by direct palmitate association with the Ptc receptor, as described above. However, another possibility is that Shh palmitoylation and the resulting membrane association of the N-terminal Shh peptide assures its proteolytic removal and thus prevents any unprocessed Shh release. Notably, unprocessed N-terminal peptides block Shh binding sites for the receptor Ptc, while shedding removes these inhibitory peptides (Ohlig et al., 2011; Schürmann et al., 2018). Consistent with this hypothesis, artificial elimination of the palmitate (in Shh^C25S^ or upon Shh expression without sufficient Hhat function) results in soluble Hhs with most of their Ptc binding sites still blocked by the uncleaved inhibitory N-peptide (Ohlig et al., 2011; Schürmann et al., 2018), and artificial elimination of the non-palmitoylated N-terminal peptide increases Shh biofunction (Jakobs et al., 2017; Ohlig et al., 2011; Ohlig et al., 2012). Therefore, our shedding model does not contradict the essential role of Shh palmitoylation for Hh activity regulation, but provides an alternative – yet indirect – explanation for it, both *in vitro* and *in vivo*. We finally note that a dispensable role of Shh lipids for signal receiving cells is supported by Ptc activity regulation as a consequence of (unlipidated) nanobody binding to the ECD1 of Ptc (Zhang et al.), a site that also represents the physiological interface for lipid-independent Shh binding (Gong et al., 2018).

## Acknowledgments

The excellent technical work of Sabine Kupich and Reiner Schulz is gratefully acknowledged. We thank Dr. C. Garbers (University of Magdeburg, Germany) for A10 knockout cells and A10 cDNA and Dr. S. Ogden (St. Jude Children’s Research Hospital, Memphis, TN, USA) for Disp cDNA.

## Author contributions

**Kristina Ehring**: Investigation, Visualization, Formal analysis, Writing-Original draft preparation. **Dominique Manikowski**: Investigation, Methodology, Visualization, Formal analysis, Writing-Reviewing and Editing. **Jurij Froese**: Investigation, Visualization, Formal analysis. **Petra Jakobs**: Investigation, Methodology. **Fabian Gude**: Investigation, Visualization. **Ursula Rescher**: Methodology, Resources. **Jonas Goretzko**: Investigation, Validation. **Kay Grobe**: Conceptualization, Supervision, Writing-Reviewing and Editing, Funding acquisition. **Competing interests**: Authors declare no competing interests. **Data and materials availability**: All data are available in the main text or the supplementary materials. Materials can be made available upon request.

## Materials and Methods

### Cell lines

Bosc23 cells and C3H10T1/2 reporter cells were cultured in DMEM supplemented with 10% fetal calf serum (FCS) and 100 μg/mL penicillin-streptomycin. Human embryonic kidney cells (HEK293T) and HEK293T A10 knockout cells (A10^−/−^) were kindly provided by Christoph Garbers (Otto-von-Guericke University Magdeburg, Germany) and grown in DMEM supplemented with 10% FCS and 100 μg/mL penicillin-streptomycin.

### Generation of Disp^−/−^ cells with CRISPR/Cas9

Disp1 knockout cells (Disp^−/−^) were generated according to manufacturer’s protocol (Dharmacon) from Bosc23 cells. The following RNAs and plasmids were used: 1) Edit-R Human DISP1 crRNA (CM-013596-02-0002), 2) Edit-R CRISPR-Cas9 Synthetic tracrRNA (U-002005-20), 3) Edit-R hCMV-PuroR-Cas9 Expression Plasmid (U-005100-120), and 4) Edit-R crRNA Non-targeting Control #1 (U-007501-01-05). Disp1 knockout was confirmed via sequencing of PCR products generated from the CRISPR-Cas9 targeted Disp DNA target site followed by the sequencing of 10 individual (cloned) PCR products. Disp1 knockout was further confirmed by immunoblotting with anti-Disp antibodies (R&D systems, AF3549). Non-targeting control (nt Ctrl) guide RNA did not change Disp DNA sequence or protein expression in control cells. Off-targets predicted by CRISPOR were analyzed by DNA sequencing and their wild-type sequence confirmed (Table S1). Independently generated nt Ctrl and Disp^−/−^ cell lines were used to confirm impaired Shh release and cholesterol quantification (shown in Fig. S5B). In all assays, nt Ctrl and Disp^−/−^ cell lines behaved like the lines presented in this work.

### Cloning of recombinant proteins

Shh expression constructs were generated from murine cDNA (NM_009170: nucleotides 1-1314, corresponding to amino acids 1-438 and ShhN: nucleotides 1-594, corresponding to amino acids 1-198) and human hedgehog acyltransferase (Hhat) cDNA (NM_018194). Both cDNAs were cloned into pIRES (Clontech) for their coupled expression from bicistronic mRNA to achieve near-quantitative Shh palmitoylation. ShhN (nucleotides 1-594, corresponding to amino acids 1-198) and Hhat were also cloned into pIRES. Shh^HA^ and ^C25S^Shh^HA^ (HA inserted between amino acids 197 and 198) were generated by site-directed mutagenesis (Stratagene). Unlipidated ^C25S^ShhN cDNA and non-palmitoylated ^C25A^Shh cDNA (amino acids 1-438) were inserted into pcDNA3.1 (Invitrogen). Primer sequences can be provided upon request. Human Scube2 constructs were a kind gift from Ruey-Bing Yang (Academia Sinica, Taiwan). Murine V5-tagged Disp^tg^ and V5-tagged cleavage-site deficient Disp^ΔCS^ were a kind gift from Stacey Ogden (St. Jude Children’s Research Hospital, Memphis, USA). Murine Disp^ΔL2^ was generated from Disp^tg^ by deletion of the second extracellular loop (L2) between transmembrane domain 7 and 8 (amino acids 752-972). Murine Ptc^ΔL2^ was generated from Ptc Full Length (pcDNA-h-mmPtch1-FL, Addgene #120889) by deletion of the second extracellular loop (L2) between transmembrane domain 7 and 8 (amino acids 794-997). Primer sequences can be provided upon request. Human A10 cDNA was kindly provided by Christoph Garbers (Otto-von-Guericke University Magdeburg, Germany). The A10^C^ construct was generated by fusion of the soluble A10 fragment (nucleotides 1-2001, corresponding to amino acids 1-667) with ShhC (nucleotides 580-1314, corresponding to amino acids 194-438), which results in a cholesteroylated A10 protein after autocatalytic cholesteroylation.

### Protein detection

HEK293T cells or Bosc23 cells were seeded into 6-well plates and transfected with 0.5 μg Disp, Ptc, and A10 encoding constructs or 1 μg Shh constructs with or without 0.5 μg Scube2 by using Polyfect (Qiagen). Cells were grown for 2 days or 3 days for Disp^−/−^ rescue experiments at 37°C with 5% CO_2_ in DMEM containing 10% fetal calf serum and penicillin-streptomycin (100 μg/mL). Media were changed to serum-free DMEM for 6 h, harvested, and centrifuged at 300*g* for 10 min to remove debris. Supernatants incubated with 10% trichloroacetic acid (TCA) for 30 min on ice, followed by centrifugation at 13,000*g* for 20 min to precipitate the proteins. Cell lysates and corresponding supernatants were analyzed on the same reducing SDS polyacrylamide gel and detected by Western blot analysis by using goat-α-Shh antibodies (R&D Systems, AF464), rabbit-α-GAPDH antibodies (Cell Signaling, GAPDH 14C10, #2118), or anti-β-actin antibodies (Sigma-Aldrich, A3854) followed by incubation with horseradish peroxidase-conjugated secondary antibodies. GAPDH, β-actin (for cell lysates), or Ponceau S (for media) served as loading controls. Shh release was quantified by ImageJ and calculated as the ratio of total or processed (truncated) soluble Shh relative to the cellular Shh material. Relative Shh release from control cells (nt Ctrl) was set to 100% and Shh release from Disp^−/−^ cells was expressed relative to that value. For rescue experiments, Shh release from empty vector transfected control cells was set to 100%. In cholesterol depletion experiments, serum-free medium was supplemented with 0-800 μg/mL methyl-β-cyclodextrin (MβCD) for 6h prior to TCA precipitation and subsequent immunoblotting analysis, and Shh release of mock treated cells was set to 100%. To quantify A10 dependent release, we set Shh release from HEK WT controls to 100%, and Shh release from A10^−/−^ cells was expressed relative to this value. Anti-HMGCR signals in cell lysates were first normalized against protein loading controls and values were then calculated relative to HMGCR amounts in nt Ctrl cells (set to 100%).

### Shh release in the presence of high-density lipoprotein (HDL)

nt Ctrl or Disp^−/−^ cells were transfected with pIRES for coupled Shh and Hhat expression together with Scube2 cDNA as described earlier. Two days after transfection, cells were washed twice with serum-free DMEM and additionally incubated for 1 h in serum-free DMEM. This extensive washing was intended to quantitatively remove serum lipoproteins. Serum-free DMEM was then discarded and cells were incubated in serum-free DMEM containing 120 μg/mL human HDL (Sigma-Aldrich, L1567) for 6 h. For cell debris removal, supernatants were centrifuged for 10min at 300*g*. For subsequent Shh purification, supernatants were incubated with 5 μg/mL anti-Shh antibody DSHB 5E1 for 2 h at 4°C, followed by the addition of 5 mg protein A beads (Sigma, P1406) in phosphate-buffered saline (PBS) and incubated at 4°C overnight. Immunoprecipitates were collected by centrifugation at 300*g* for 5 min and subjected to reducing SDS-PAGE followed by immunoblot analysis. Shh release was quantified by determining the ratios of soluble Shh signals detected in 5E1-Protein A pulldown samples relative to cellular actin signals. Shh release from mock-treated nt Ctrl cells (no HDL) were set to 100%.

### Shh bioactivity assay

Bosc23 cells, nt Ctrl, or Disp^−/−^ cells were transfected with Shh/Hhat or its variants thereof together with Scube2 as described earlier. Two days later, media were replaced with serum-free medium for 6 h. Media were then harvested, cellular debris removed, FCS added ad 10%, mixed 1:1 with DMEM supplemented with 10% FCS and 100 μg/mL antibiotics, and the mixture added to C3H10 T1/2 cells. Cells were lysed 6 days after osteoblast differentiation was induced, lysed in 1% TritonX-100 in PBS, and osteoblast-specific alkaline phosphatase activity measured at 405 nm by using 120 mM p-nitrophenolphosphate (Sigma) in 0.1 M Tris buffer (pH 8.5). Mock-treated C3H10 T1/2 cells served as negative controls.

### Total (esterified) and free (unesterified) cholesterol quantification

To quantify the content of total and free cholesterol in nt Ctrl cells and Disp^−/−^ cells, we seeded cells into 6-well plates and grew the cells in DMEM containing 10% FCS and 100 μg/mL penicillin-streptomycin at 37°C for 2 days. For experiments containing MβCD, cells were incubated with 800 μg/mL MβCD for 6 h in serum-free media prior to cell lysis. Afterwards, cells were washed twice with PBS, harvested by using cell scrapers and 1 mL PBS, centrifuged at 800*g* for 5 min at 4°C and cell pellets resuspended in 500 μL HB buffer (250 mM sucrose, 3 mM imidazole at pH 7.4) for washing. Samples were centrifuged again and cell pellets were resuspended in 400 μL HB buffer. Cells were subsequently mechanically lysed by using a syringe (27G) and nuclei removed by centrifugation at 1000*g* for 15 min at 4°C. Total protein concentration of supernatants were measured at 280 nm by using a NanoDrop spectrophotometer and adjusted to similar levels. For total and free cholesterol quantification, Amplex Red cholesterol assay (Invitrogen, A12216) was conducted according to the manufacturer’s protocol with or without the use of the cholesterol esterase (converts esterified cholesteryl into free cholesterol), respectively.

### Quantitative real-time PCR

To determine the relative messenger RNA (mRNA) expression changes of genes involved in cholesterol homeostasis, we performed a qPCR assay. RNA was extracted from Disp−/− and nt Ctrl cells with TRIzol reagent (Invitrogen), and equal RNA amounts were reverse-transcribed for cDNA synthesis by using the first-strand DNA synthesis kit (ThermoFisher) according to the manufacturer’s instructions. mRNA levels were determined by using Rotor-Gene SYBR Green (Qiagen, 204074) on a BioRad CFX384 Real-Time-System C1000 Touch Thermal Cycler. The qPCR primers are listed in Table S2. β-actin was used as a reference gene and the log2-fold change was calculated with the 2−ΔΔct method.

### Confocal microscopy

Disp^−/−^ cells were seeded onto gelatin-coated 4-well cell culture chamber slides (PAA, PAA30104X) and transfected with either Shh/Hhat and Scube2 or V5-tagged Disp^ΔCS^ together with empty eGFP-N1 vectors (for cytoplasm visualization) using Polyfect as described earlier. After 2 days in culture, cells were fixed with 4% PFA for 10 min at room temperature under non-permeabilizing conditions. Mouse-α-Shh (DSHB, 5E1) and mouse-α-V5 antibodies (Abcam, ab27671) were used to stain Shh and Disp, respectively. Texas Red-conjugated goat-α-mouse IgG antibodies and a Zeiss LSM700 confocal microscope were used for visualization. DAPI was used as a nuclear counterstain. Representative confocal planes are shown.

### Reverse-phase high performance liquid chromatography (HPLC)

Bosc23 cells were transfected with expression plasmids for unlipidated ^C25A^ShhN control protein and cholesteroylated (yet non-palmitoylated) ^C25A^Shh. Two days after transfection, cells were lysed in radioimmunoprecipitation assay buffer containing complete protease inhibitor cocktail (Roche, Basel, Switzerland) on ice and ultracentrifuged, and the soluble whole-cell extract was acetone precipitated. Protein precipitates were resuspended in 35 μL of (1,1,1,3,3,3) hexafluoro-2-propanol and solubilized with 70 μL of 70% formic acid, followed by sonication. Reverse-phase HPLC was performed on a C4-300 column (Tosoh, Tokyo, Japan) and an Äkta Basic P900 Protein Purifier. To elute the samples, we used a 0-70% acetonitrile/water gradient with 0.1% trifluoroacetic acid at room temperature for 30 min. Eluted samples were vacuum dried, resolubilized in reducing sample buffer, and analyzed by SDS-PAGE and immunoblotting by using anti-Shh antibodies (R&D Systems, AF464). Signals were quantified with ImageJ and normalized to the highest protein amount detected in each run.

### Bioanalytical and statistical analysis

All statistical analyses were performed in GraphPad Prism. Applied statistical tests, post hoc tests, and number of independently performed experiments are stated in the figure legends. A p-value of < 0.05 was considered statistically significant. *: p<0.05, **: p<0.01, ***: p<0.001, and ****: p<0.0001 in all assays. Error bars represent the standard deviations of the means.

## Supplemental Data

**Fig. S1.**
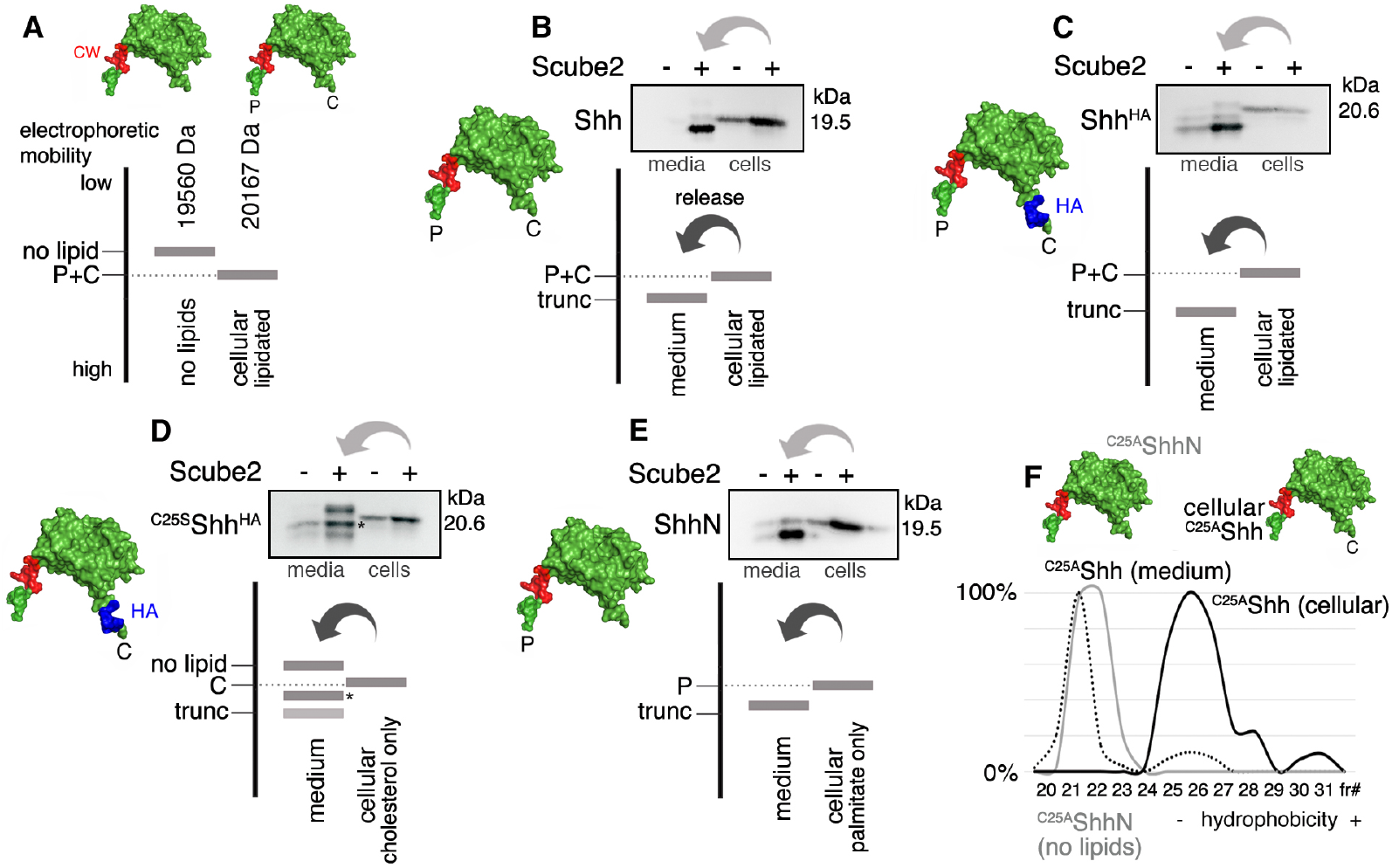
Schematic representation of proteolytic Shh processing. Schematics of expressed Shh constructs (before their release) are shown. **A)** As previously published (Pepinsky et al., 1998; Porter et al., 1996), dual-lipidated Hh (cellular lipidated) migrates faster in SDS-PAGE than does *E. coli*-expressed unlipidated Hh (no lipids), although mass spectrometry determined molecular weights of 20167 Da for the former form and 19560 Da for the latter. Increased electrophoretic cellular lipidated Shh mobility, despite its higher molecular weight, was explained by SDS association with the large hydrophobic sterol backbone of cholesterol and the C_16_ hydrocarbon tail of the palmitate. Consistent with this, chemical hydrolysis of the ester bond that attaches cholesterol to Shh was described to decrease electrophoretic mobility of the delipidated product (Zeng et al., 2001). **B)** Increased electrophoretic mobility of soluble Shh over the dual-lipidated cellular protein (P+C) results from the loss of both lipidated terminal peptides during release: Shh delipidation decreases its electrophoretic mobility (see A), but the additional loss of associated terminal peptides (trunc) compensates for this decrease. As a consequence, proteolytic Shh processing results in a small net increase in electrophoretic mobility. Note that coupled Shh release and processing depends on Scube2. Bottom: schematic of Shh release. **C)** Insertion of a C-terminal HA tag supports this hypothesis: removal of terminal peptides, including the 1 kDa tag, increases the net electrophoretic mobility gain of solubilized proteins compared with the dual-lipidated (tagged) cellular precursor (Jakobs et al., 2014; Jakobs et al., 2017). **D)** Genetic deletion of the N-terminal palmitate (C25S) impairs ^C25S^Shh^HA^ conversion into the fully processed protein. Instead, a C-processed but N-terminally unprocessed intermediate (asterisk) is released (Jakobs et al., 2014; Jakobs et al., 2017), suggesting that N-palmitate (by its continued membrane association) controls quantitative Shh N-processing (Ohlig et al., 2011). An additional decreased electrophoretic mobility fraction that still carries the C-terminal HA tag (top band, compare with A) is also detected, suggesting generation of a second cleavage site proximal to the cholesterol as a consequence of HA insertion. **E)** ShhN co-expression with Hhat (resulting in N-palmitoylated proteins lacking the cholesterol moiety) also results in the solubilization of truncated proteins (compare with Scube2 independent secretion of non-palmitoylated ^C25S^ShhN, Fig. 1C). **F)** Reverse-phase HPLC confirms Shh delipidation during release. Non-lipidated soluble control ^C25A^ShhN (gray solid line) and cholesterol-modified ^C25A^Shh (dotted line) were released from Disp-expressing Bosc23 cells in the presence of Scube2. Both soluble proteins bound to and eluted from a hydrophobic C4 column in a similar manner, but soluble ^C25A^Shh was less hydrophobic than its cholesteroylated precursor in Bosc23 cell lysates (black solid line). This demonstrates loss of the cholesteroylated C-terminus during release. Elution profiles are expressed relative to the highest protein amount in a given fraction (set to 100%). fr#: fraction number.

**Fig. S2.**
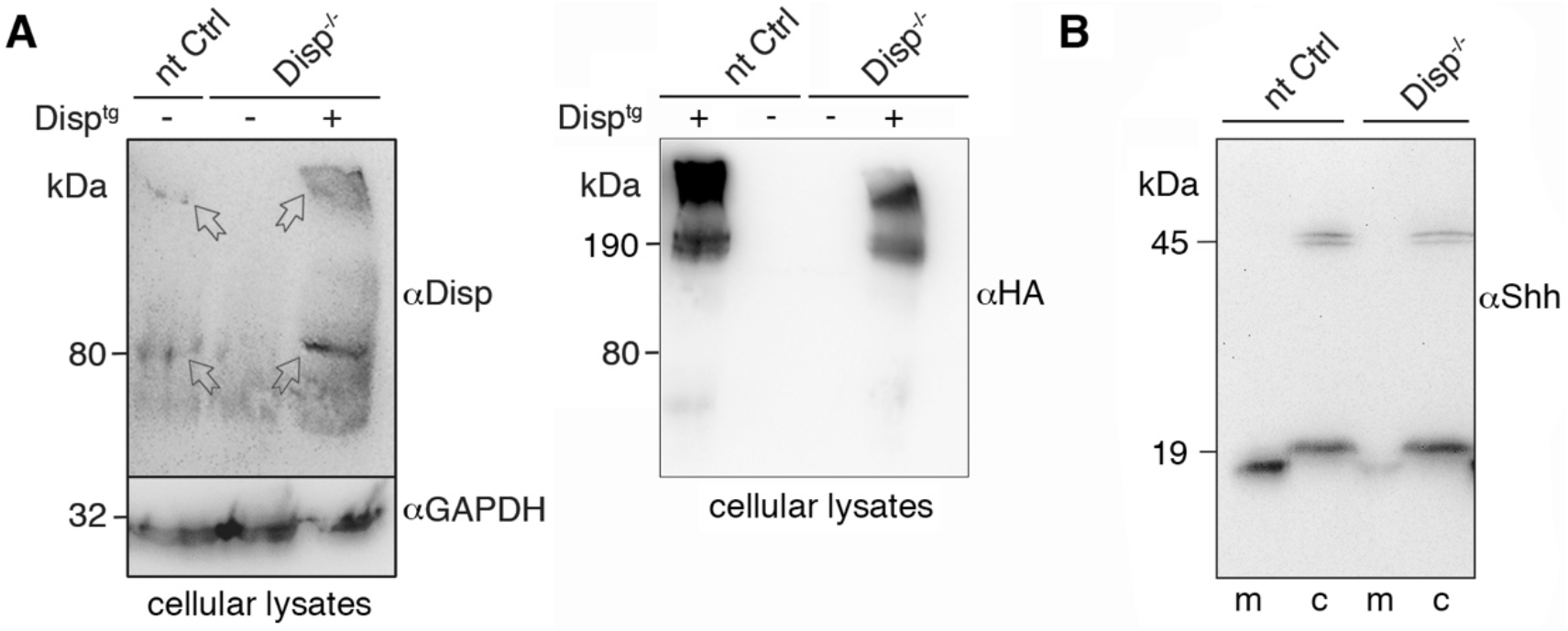
Loss of Disp protein expression and unimpaired Shh autoprocessing in Disp^−/−^ cells. **A)** Left: Disp^−/−^ cellular lysates were subjected to SDS-PAGE/immunoblotting. Arrows indicate full-length Disp protein and 80 kDa Disp degradation products in nt Ctrl and Disp^tg^-expressing Disp^−/−^ cells. Right: The established large size of Disp clusters not disrupted by SDS-PAGE (Stewart et al., 2018) was confirmed by specific detection of HA-tagged Disp^tg^. **B)** Similar autoprocessing of 45 kDa Shh precursor proteins into cholesteroylated, 19 kDa products in nt Ctrl and Disp^−/−^ cells (c).

**Fig. S3.**
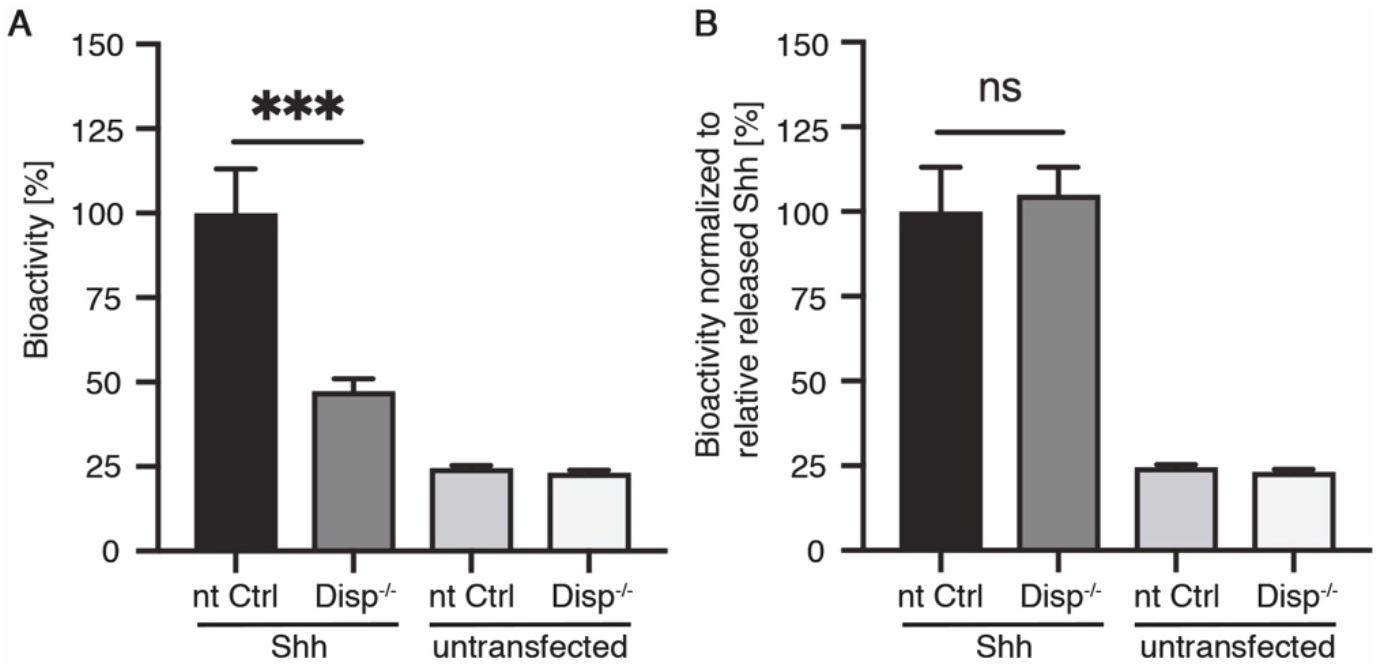
Soluble truncated Shh is bioactive. **A)** Quantification of Shh bioactivity. Shh was released from transfected and untransfected nt Ctrl and Disp^−/−^ cells and the conditioned medium subsequently adjusted to 10% FCS and added to C3H10T1/2 reporter cells. Compared with Shh release from nt Ctrl cells, reduced Shh solubilization from Disp^−/−^ cells resulted in reduced Shh-dependent differentiation of C3H10T1/2 cells into alkaline phosphatase-producing osteoblasts. n=3 datasets from 1 experiment, one-way ANOVA, Sidak’s multiple comparison post hoc test. **B)** Normalization of soluble protein amounts demonstrates that Shh released from Disp^−/−^ cells is bioactive. This also suggests that reduced Shh biofunction as a consequence of Disp deletion results from its reduced solubilization and not its processing (e.g. the associated loss of the N-palmitate). n=3 datasets obtained from 1 experiment, one-way ANOVA, Sidak’s multiple comparison post hoc test.

**Fig. S4.**
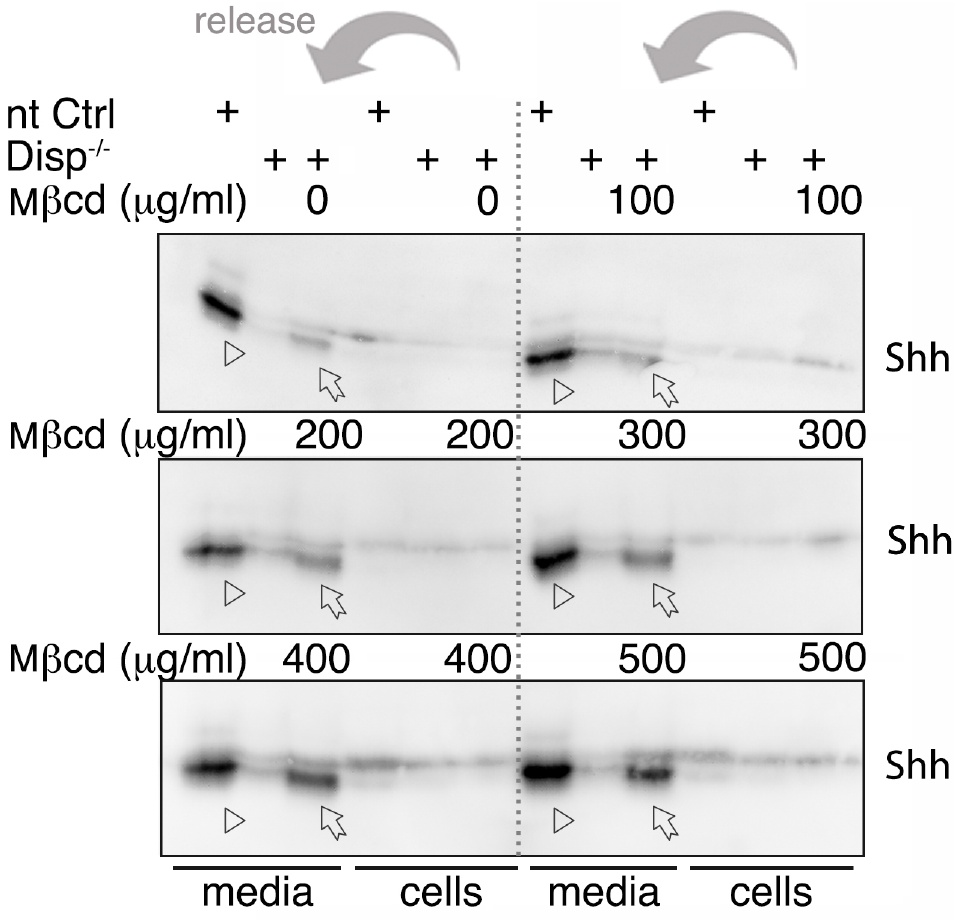
MβCD restores Shh release from Disp^−/−^ cells in a dose-dependent manner. 0 to 500 μg/mL MβCD was used to increase processed Shh release from Disp^−/−^ cells in the respective lane (arrows) compared with untreated Disp^−/−^ cells (adjacent lane). Reference Shh amounts released from nt Ctrl cells in the absence of MβCD are also shown (arrowheads).

**Fig. S5.**
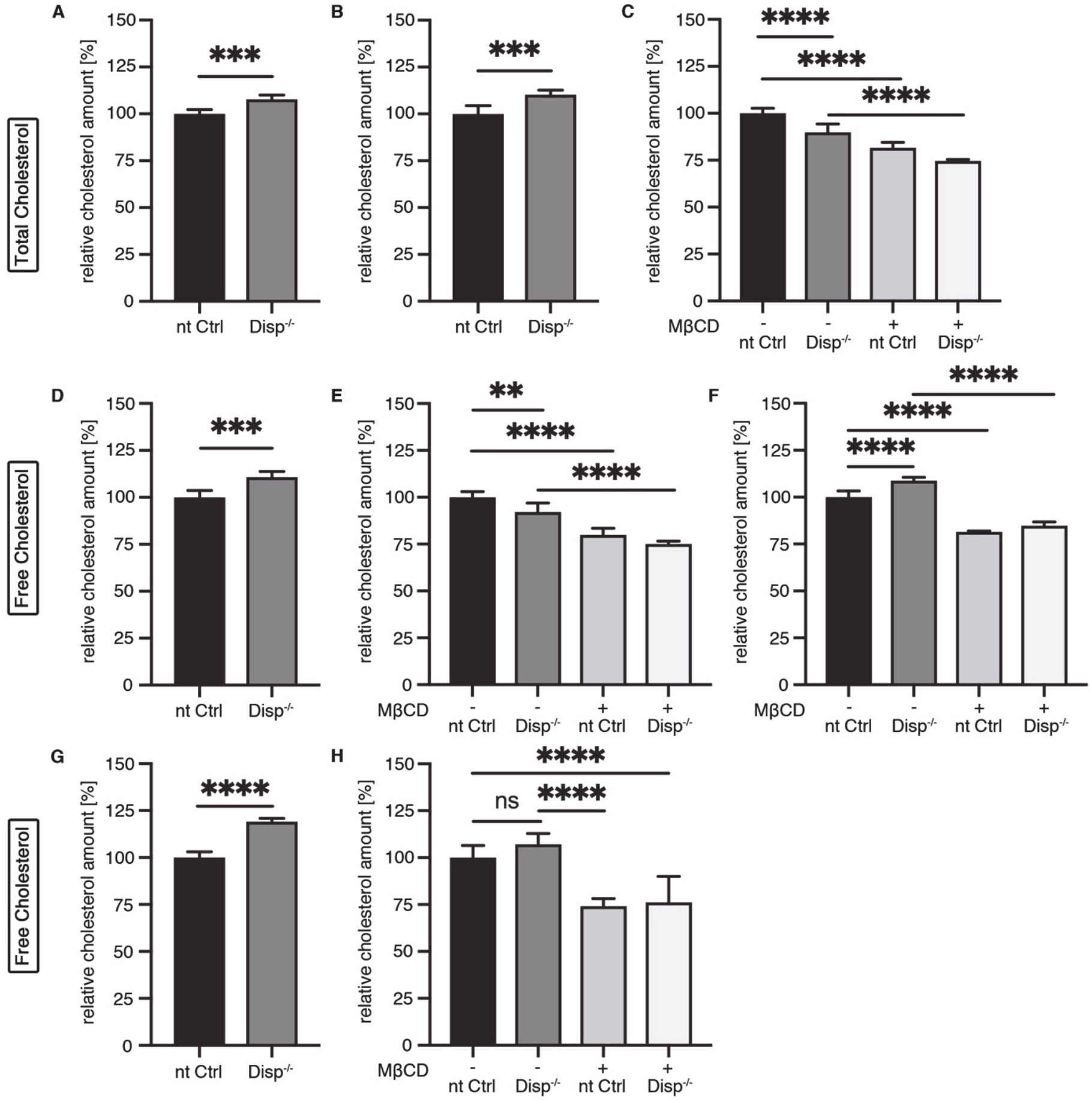
Quantification of total and free (unesterified) cholesterol in nt Ctrl and Disp^−/−^ cells. Eight assays were independently conducted. **A-H)** Two of 3 total cholesterol quantification assays (A, B) and 3 of 5 free cholesterol quantification assays (D, F, G) showed slightly increased cholesterol contents in Disp^−/−^ cells compared with nt Ctrl cells. However, 2 of the remaining assays (C, E) showed the opposite effect and the assay in H did not show a significant difference between nt Ctrl and Disp^−/−^ cells. The averaged values of D-H are summarized in Fig. 4E. MβCD consistently depleted free cellular cholesterol, as reflected by decreased total and free cholesterol (C, E, F, H). A: n=5 datasets from 1 experiment. B: A second independent nt Ctrl and Disp^−/−^ cell line displayed similar results, n=6 datasets from 1 experiment. C: n=6 datasets from 1 experiment. D: n=6 datasets from 1 experiment. E: n=6 datasets from 1 experiment. F: n=6 datasets from 1 experiment. G: n=5 datasets from 1 experiment. H: n=6 datasets from 1 experiment. A, B, D, G: Unpaired t-test in each assay. C, E, F, H: One-way ANOVA, Sidak’s multiple comparison post hoc test in each assay.

**Fig. S6.**
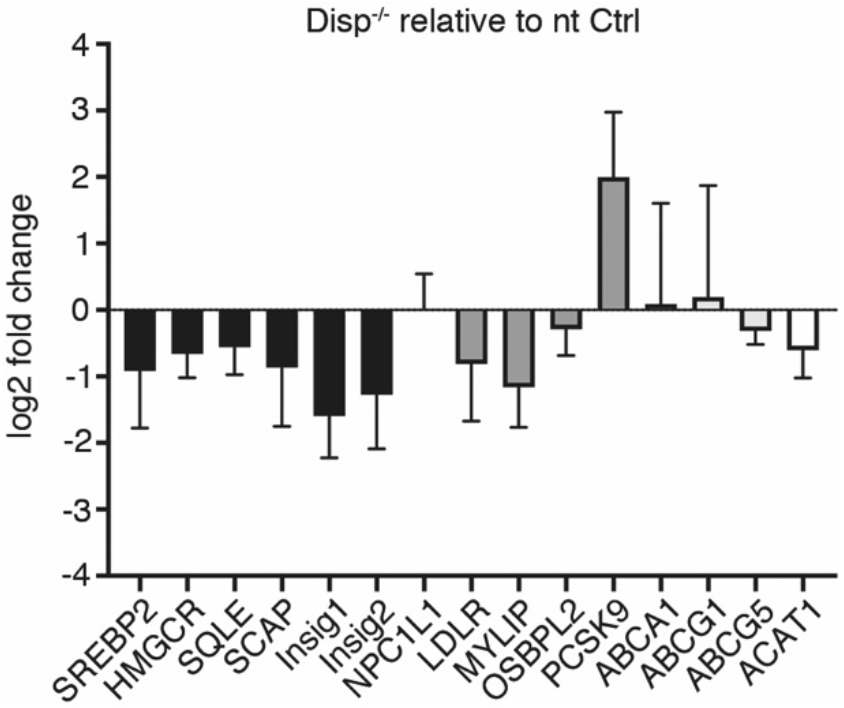
qPCR did not reveal strong changes in mRNA expression of key proteins involved in cellular cholesterol homeostasis. **Black** bars represent genes involved in cholesterol biosynthesis: sterol regulatory element-binding protein 2 (*SREBP2*) – a transcriptional regulator of cholesterol biosynthesis; HMG-CoA-reductase (*HMGCR*) – a rate-limiting enzyme in cholesterol biosynthesis; squalene monooxygenase (*SQLE*) – another rate-limiting enzyme in cholesterol biosynthesis; sterol regulatory element-binding protein cleavage-activation protein (*SCAP*) – a sensor of free cholesterol; Insulin-induced gene 1 protein (*Insig1*) – a feedback regulator of cholesterol synthesis; Insulin-induced gene 2 protein (*Insig2*) – feedback control of cholesterol synthesis. The **dark gray** bar represents a gene involved in the uptake of cholesterol: NPC1-like intracellular cholesterol transporter 1 (*NPC1L1*) – active uptake of dietary cholesterol. **Gray** bars represent genes involved in cellular cholesterol uptake: low-density lipoprotein receptor (*LDLR*) – receptor for cholesterol-rich LDL; E3 ubiquitin-protein ligase (*MYLIP*) – mediates ubiquitination and degradation of the LDLR; oxysterol-binding protein-related protein 2 (*OSBPL2*) – intracellular transport protein for sterols and phospholipids; proprotein convertase subtilisin/kexin type 9 (*PCSK9*) – degrades the LDLR during biosynthesis or recycling, thereby reducing the uptake of cholesterol into the cell. **Light gray** bars represent genes involved in cellular cholesterol efflux: phospholipid-transporting ATPase (*ABCA1*) – translocation of phospholipids and cholesterol from the cytoplasmic to the extracellular leaflet and transfer to apolipoproteins; ATP-binding cassette sub-family G member 1 (ABCG1) – efflux of phospholipids such as sphingomyelin or cholesterol to HDL; ATP-binding cassette sub-family G member 5 (*ABCG5*) – mediates sterol transport across the cell membrane to HDL. The **white bar** represents a gene involved in cholesterol esterification and storage: Acetyl-CoA acetyltransferase (*ACAT1*) – formation of cholesteryl esters to prevent accumulation of unesterified cholesterol. n=2 qPCR datasets obtained from independently conducted experiments.

**Fig. S7.**
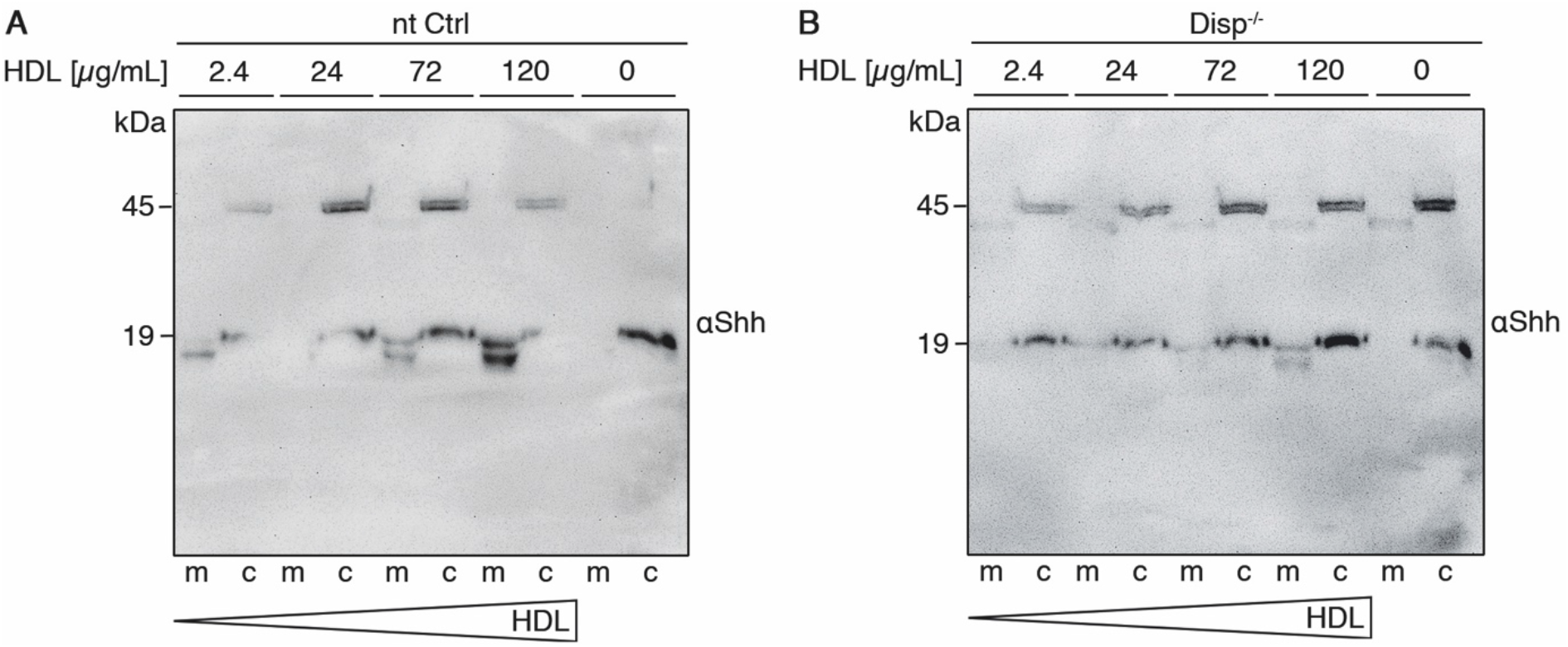
Increasing HDL dose dependently enhances Shh release in nt Ctrl cells, but much less so in Disp^−/−^ cells. **A, B)** HDL concentrations ranging from 2.4 μg/mL to 120 μg/mL increasingly facilitated Shh release in nt Ctrl cells, but less so in Disp^−/−^ cells. As described in Fig. 1, the top (seemingly unprocessed) soluble Shh likely represents the non-palmitoylated, N-terminally unprocessed but C-terminally processed protein.

**Table S1.** Predicted guide RNA off-targets are not affected. Predicted off-target sites of Disp1 guide RNA by CRISPOR were analyzed by DNA sequencing and their wild-type (WT) sequences were confirmed.

| Predicted off-target | Gene bank number | Predicted off-target site |
| --- | --- | --- |
| YLPM1 | NM_019589 | WT sequence confirmed |
| LINC01304 | NR_037881 | WT sequence confirmed |
| APLF | NM_173545 | WT sequence confirmed |
| MINK1 | NG_028005 | WT sequence confirmed |
| SLC26A9 | NM_052934 | WT sequence confirmed |

**Table S2.**
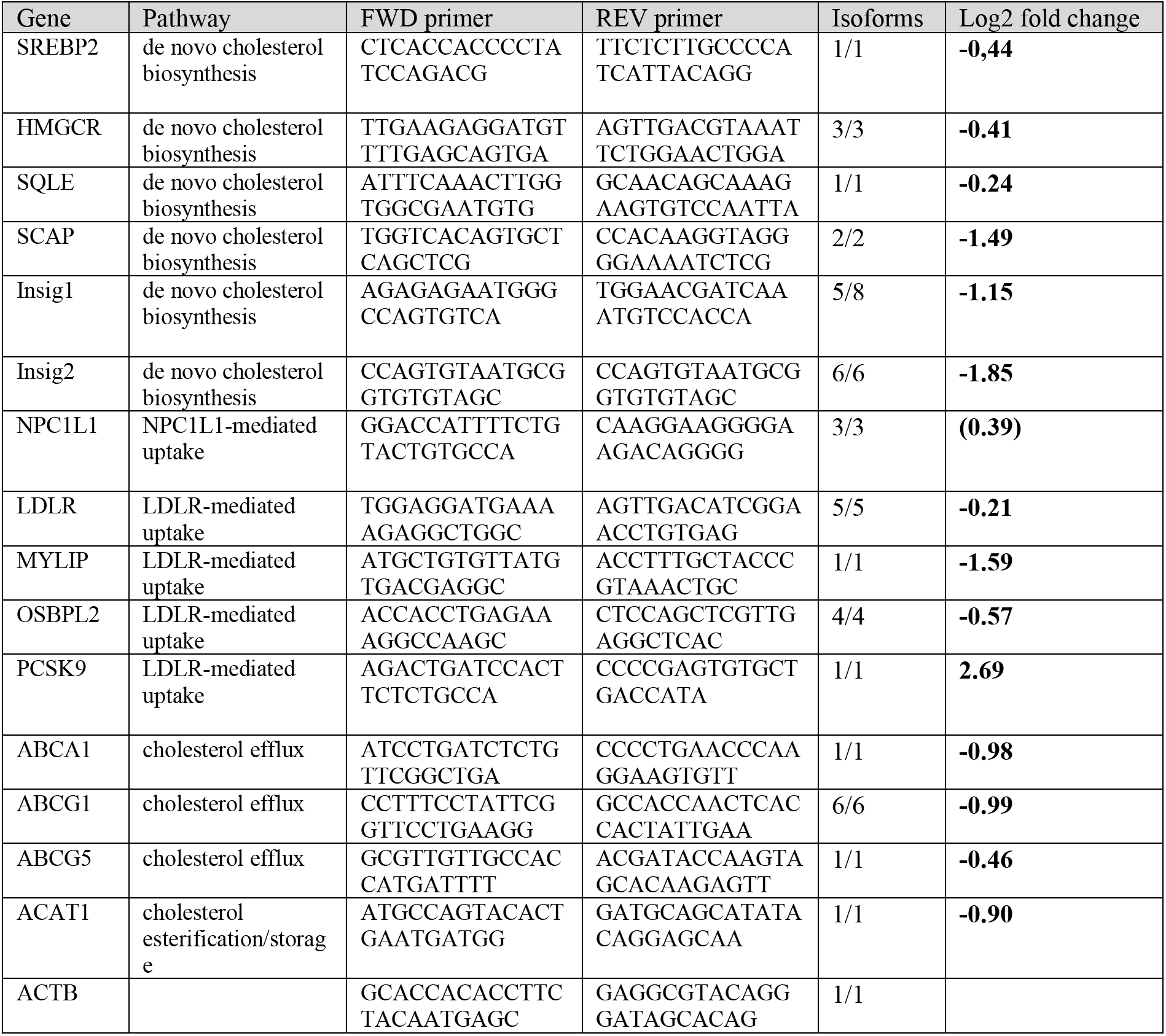
qPCR results. Genes and primers used for their detection are shown. Log2 fold change describes expression in Disp^−/−^ cells compared with nt Ctrl cells. Values <1 indicate reduced transcription in Disp^−/−^ cells; values >1 indicate a transcriptional increase.

**Table S3.** Raw data used for the figures in this study.

| Figure | Genotype | Constructs used/treatment | Values (Mean) | s.d. |
| --- | --- | --- | --- | --- |
| 1E |  | Shh | 0.698 (A<sub>405</sub>, arbitrary units) | 0.11 |
|  |  | C25S<sup>Shh</sup><sup>HA</sup> | 0.65 (A<sub>405</sub>, arbitrary units) | 0.042 |
|  |  | Shh<sup>HA</sup> | 0.62 (A<sub>405</sub>, arbitrary units) | 0.02 |
|  |  | mock | 0.35 (A<sub>405</sub>, arbitrary units) | 0.015 |
| 2B' | nt Ctrl |  | 100% | 21.07 |
|  | Disp<sup>-/-</sup> |  | 26.78% | 20.63 |
| 2C' | nt Ctrl |  | 99.99% | 29.41 |
|  | Disp<sup>-/-</sup> |  | 70.99% | 56.45 |
| 2D' | nt Ctrl |  | 100% | 8.443 |
|  | Disp<sup>-/-</sup> |  | 63.76% | 29.76 |
| 2E' | nt Ctrl |  | 99.99% | 36.86 |
|  | Disp<sup>-/-</sup> |  | 76.40% | 59.34 |
| 3A' | nt Ctrl |  | 100% | 1.838 |
|  | Disp<sup>-/-</sup> |  | 78.64% | 26.54 |
| 3B' | nt Ctrl |  | 100% | 30.24 |
|  | Disp<sup>-/-</sup> |  | 76.76% | 64.48 |
| 3D' | Disp<sup>-/-</sup> | empty vector | 100% | 35.07 |
|  | Disp<sup>-/-</sup> | Disp<sup>tg</sup> | 263.6% | 136.4 |
|  | Disp<sup>-/-</sup> | Disp<sup>ΔL2</sup> | 201.3% | 81.26 |
| 3E' | nt Ctrl | empty vector | 100% | 32.84 |
|  | nt Ctrl | Disp<sup>tg</sup> | 146.4% | 115.1 |
|  | nt Ctrl | Disp<sup>ΔL2</sup> | 138.9% | 59.66 |
| 4B' | Disp<sup>-/-</sup> | empty vector | 100% | 30.89 |
|  | Disp<sup>-/-</sup> | Ptc<sup>tg</sup> | 320.1% | 119.0 |
|  | Disp<sup>-/-</sup> | Ptc<sup>ΔL2</sup> | 280.7% | 114.2 |
| 4C' | nt Ctrl | empty vector | 100% | 35.21 |
|  | nt Ctrl | Ptc<sup>tg</sup> | 156.5% | 39.84 |
|  | nt Ctrl | Ptc<sup>ΔL2</sup> | 149.9% | 61.50 |
| 4D' | Disp<sup>-/-</sup> | mock | 100% | 31.45 |
|  | Disp<sup>-/-</sup> | MβCD (800 μg/mL) | 505.2% | 168.8 |
| 4E | nt Ctrl | mock | 100% | 3.863 |
|  | Disp<sup>-/-</sup> | mock | 107.2% | 9.505 |
|  | nt Ctrl | MβCD (800 μg/mL) | 77.91% | 4.628 |
|  | Disp<sup>-/-</sup> | MβCD (800 μg/mL) | 77.48% | 9.175 |
| 4F' | nt Ctrl |  | 100% | 8.159 |
|  | Disp<sup>-/-</sup> |  | 83.50% | 36.16 |
| 4G' | nt Ctrl | mock | 100% | 2.530 |
|  | nt Ctrl | HDL (120 μg/mL) | 178.3% | 41.49 |
|  | Disp<sup>-/-</sup> | mock | 18.80% | 9.783 |
|  | Disp<sup>-/-</sup> | HDL (120 µg/mL) | 62.00% | 19.01 |
| 5A' | WT |  | 100% | 0 |
|  | A10<sup>-/-</sup> |  | 35.99% | 36.31 |
| 5C' | WT | empty vector | 100% | 0 |
|  | A10<sup>-/-</sup> | empty vector | 10.22% | 15.29 |
|  | A10<sup>-/-</sup> | Disp<sup>tg</sup> | 28.65 | 14.45 |
| 5D' | Disp<sup>-/-</sup> | empty vector | 100% | 0.1 |
|  | Disp<sup>-/-</sup> | A10<sup>tg</sup> | 152.5% | 30.65 |
|  | Disp<sup>-/-</sup> | A10<sup>c</sup> | 227.7% | 46.32 |
| S3A | nt Ctrl | Shh | 100% | 13.05 |
|  | Disp<sup>-/-</sup> | Shh | 47.32% | 3.666 |
|  | nt Ctrl | mock | 24.61% | 0.7778 |
|  | Disp<sup>-/-</sup> | mock | 23.22% | 0.6959 |
| S3B | nt Ctrl | Shh | 100% | 13.05 |
|  | Disp<sup>-/-</sup> | Shh | 105% | 8.136 |
|  | nt Ctrl | mock | 24.61% | 0.7778 |
|  | Disp<sup>-/-</sup> | mock | 23.22% | 0.6959 |
| S5A | nt Ctrl |  | 100% | 2.254 |
|  | Disp<sup>-/-</sup> |  | 107.8% | 2.269 |
| S5B | nt Ctrl |  | 100% | 4.482 |
|  | Disp<sup>-/-</sup> |  | 110.3% | 2.442 |
| S5C | nt Ctrl | mock | 100% | 2.733 |
|  | Disp<sup>-/-</sup> | mock | 89.84% | 4.419 |
|  | nt Ctrl | MβCD (800 µg/mL) | 81.56% | 2.925 |
|  | Disp<sup>-/-</sup> | MβCD (800 µg/mL) | 74.65% | 0.6654 |
| S5D | nt Ctrl |  | 100% | 3.662 |
|  | Disp<sup>-/-</sup> |  | 110.8% | 3.165 |
| S5E | nt Ctrl | mock | 100% | 3.126 |
|  | Disp<sup>-/-</sup> | mock | 92.09% | 4.913 |
|  | nt Ctrl | MβCD (800 µg/mL) | 79.96% | 3.516 |
|  | Disp<sup>-/-</sup> | MβCD (800 µg/mL) | 75.15% | 1.441 |
| S5F | nt Ctrl | mock | 100% | 3.349 |
|  | Disp<sup>-/-</sup> | mock | 108.8% | 1.801 |
|  | nt Ctrl | MβCD (800 µg/mL) | 81.51% | 0.4488 |
|  | Disp<sup>-/-</sup> | MβCD (800 µg/mL) | 84.74% | 2.085 |
| S5G | nt Ctrl |  | 100% | 3.108 |
|  | Disp<sup>-/-</sup> |  | 119.1% | 1.800 |
| S5H | nt Ctrl | mock | 100% | 6.439 |
|  | Disp<sup>-/-</sup> | mock | 107.2% | 5.698 |
|  | nt Ctrl | MβCD (800 µg/mL) | 74.06 | 4.110 |
|  | Disp<sup>-/-</sup> | MβCD (800 µg/mL) | 76.18 | 13.85 |
| S6 |  | SREBP2 | -0.9200 (log2 fold change) | 0.8575 |
|  |  | HMGCR | -0.6633 (log2 fold change) | 0.3557 |
|  |  | SQLE | -0.5600 (log2 fold change) | 0.4158 |
|  |  | SCAP | -0.8650 (log2 fold change) | 0.8839 |
|  |  | Insig1 | -1.595 (log2 fold change) | 0.6293 |
|  |  | Insig2 | -1.275 (log2 fold change) | 0.8132 |
|  |  | NPC1L1 | 0.02000 (log2 fold change) | 0.5233 |
|  |  | LDLR | -0.8150 (log2 fold change) | 0.8556 |
|  |  | MYLIP | -1.165 (log2 fold change) | 0.6010 |
|  |  | OSBPL2 | -0.2900 (log2 fold change) | 0.3960 |
|  |  | PCSK9 | 2.000 (log2 fold change) | 0.9758 |
|  |  | ABCA1 | 0.09000 (log2 fold change) | 1.513 |
|  |  | ABCG1 | 0.1950 (log2 fold change) | 1.676 |
|  |  | ABCG5 | -0.3150 (log2 fold change) | 0.2051 |
|  |  | ACAT1 | -0.6050 (log2 fold change) | 0.4172 |

